# Structural Development of Speech Networks in Young Children

**DOI:** 10.1101/2024.08.23.609470

**Authors:** Marilyn Curtis, Mohammadreza Bayat, Dea Garic, Alliete R. Alfano, Melissa Hernandez, Madeline Curzon, Andrea Bejarano, Pascale Tremblay, Paulo Graziano, Anthony Steven Dick

**Author notes:** **Correspondence:** Anthony Steven Dick PhD, Department of Psychology and the Center for Children and Families, Florida International University, Miami, Florida, 33199, United States.

## Abstract

Characterizing the structural development of the neural speech network in early childhood is important to understand speech acquisition. To investigate speech in the developing brain, 94 children aged 4-7-years-old were scanned using diffusion weighted imaging (DWI) magnetic resonance imaging (MRI). In order to increase sample size and performance variability, we included children who were diagnosed with attention-deficit hyperactivity disorder (ADHD) from a larger ongoing study. Additionally, each child completed the Syllable Repetition Task (SRT), a validated measure of phoneme articulation. The DWI data were modeled using restriction spectrum imaging (RSI) to measure restricted and hindered diffusion properties in both grey and white matter. Consequently, we analyzed the diffusion data using both whole brain analysis, and automated fiber quantification (AFQ) analysis to establish tract profiles for each of six fiber pathways thought to be important for supporting speech development. In the whole brain analysis, we found that SRT performance was associated with restricted diffusion in bilateral inferior frontal gyrus, *pars opercularis*, right pre-supplementary and supplementary motor area, and bilateral cerebellar grey matter (*p* < .005). Age moderated these associations in left *pars opercularis* and frontal aslant tract (FAT). However, in both cases only the cerebellar findings survived a cluster correction. We also found associations between SRT performance and restricted diffusion in cortical association fiber pathways, especially left FAT, and in the cerebellar peduncles. Analyses using automated fiber quantification (AFQ) highlighted differences in high and low performing children along specific tract profiles, most notably in left but not right FAT, in bilateral SLFIII, and in the cerebellar peduncles. These findings suggest that individual differences in speech performance are reflected in structural grey and white matter differences as measured by restricted and hindered diffusion metrics, and offer important insights into developing brain networks supporting speech in very young children.

## 1| INTRODUCTION

Contemporary neurobiological models of speech have clarified the primary brain regions and connections that are putatively important for speech. Such models developed in adults are potentially relevant for understanding developing speech in young children, an understudied population with regard to the neurobiology of speech. In fact, very little is known about the development of neural speech pathways in early childhood (i.e., 4- to 8-years; although see (Broce et al., 2015; S.-E. Chang et al., 2015; Johnson et al., 2022). This highlights a major deficit in our knowledge of the neurobiology of speech development. The neurobiological models based in adults allow for some expectation about which brain regions and fiber pathways might be sensitive to individual differences in speech ability as children develop in this age range, but this remains untested. With an understanding of how speech develops in the brain, we can build better developmental models relevant for early disorders of speech production, which is the focus of the present study. In the following sections, we review the relevant brain regions and fiber pathways proposed to support speech production in the developing brain, the novel diffusion-weighted imaging measures we employ to index this development, and the age-appropriate task we use to explore behavioral associations with speech production performance in young children, including those with risk for speech production difficulties such as children with attention-deficit-hyperactivity-disorder (ADHD).

### 1.1| Brain regions and fiber pathways supporting speech production

The two pertinent contemporary models of speech neurobiology that will frame this investigation–Guenther’s GO-DIVA model (Guenther, 2016) and Hickok’s updated model (Hickok et al., 2023) –describe how speech can be construed as a broad, distributed network that includes subcortical and cortical structures, and the white matter pathways which connect them (see Figure 1 for an overview of relevant structures and pathways). According to Guenther, the neural speech network can be parsed into bilateral cortical and subcortical information loops that shape the motor cortical commands for speech (Guenther, 2016). The cortico-basal ganglia loop connects basal ganglia structures (globus pallidus, substantia nigra, caudate, and the putamen), and the ventral lateral nucleus of the thalamus, with various regions of the cerebral cortex, such as the supplementary motor area (SMA) and pre-SMA, the primary motor cortex, the somatosensory cortex, and the premotor cortex (Guenther, 2016). The information loop connecting these subcortical structures plays a key role in the process of selecting and initiating appropriate motor programs for speech (Guenther, 2016). The cortico-cerebellar loop, consisting of projections through the pons, deep cerebellar nuclei, the thalamus, and the cerebral cortex, is crucial in the generation of finely timed muscle activations necessary for rapid speech production (Guenther, 2016). The information loops described provide a direct pathway connecting the subcortical structures with the cerebral cortex, where somatosensory and motor representations are integrated, and the motor commands necessary for speech production are generated and initiated.

**FIGURE 1.**
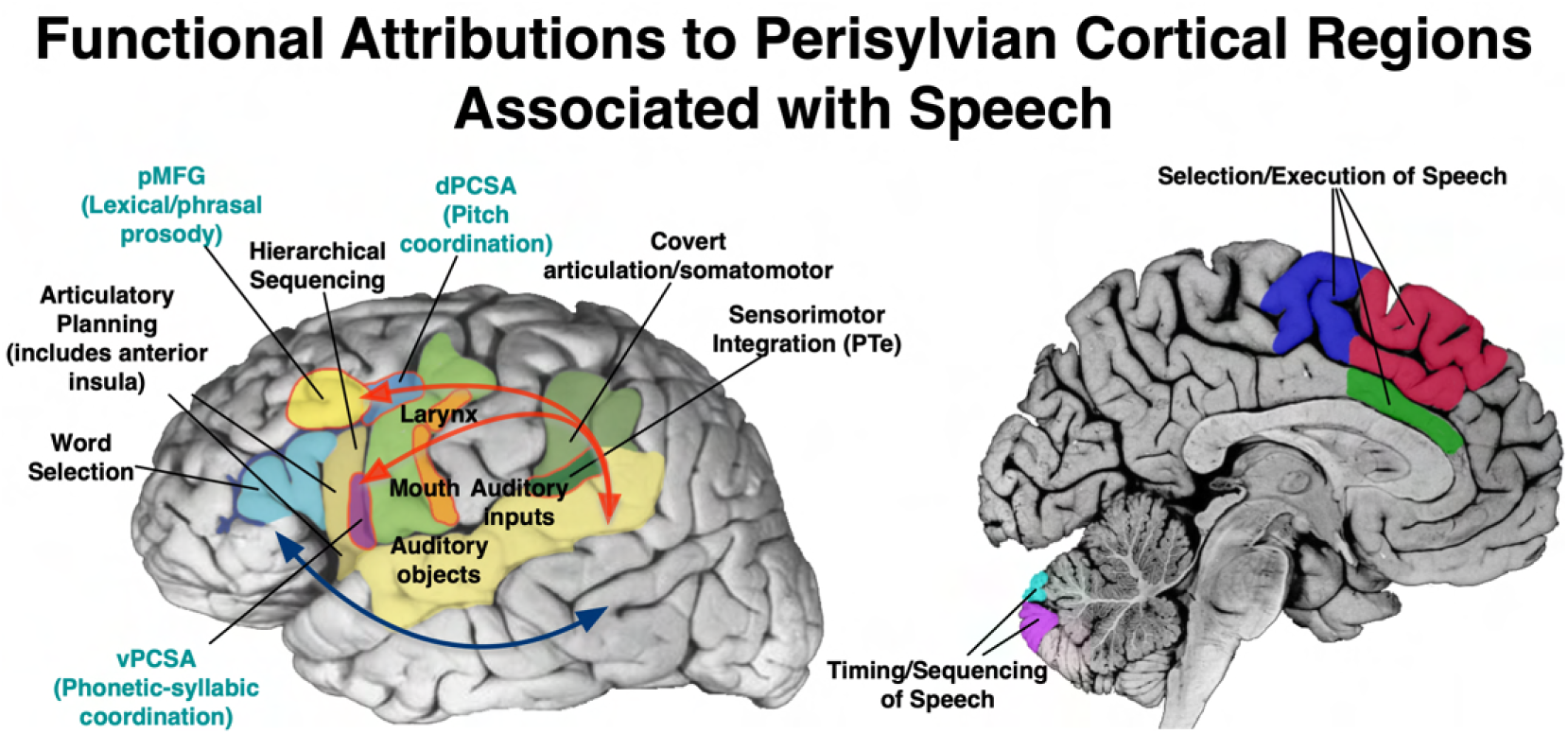
Cortical regions supporting speech are examined in the current study, situated within a broader bilateral but strongly left-lateralized perisylvian language network. The dorsal stream (upper red arrows) is primarily proposed to support sensorimotor speech processing via posterior superior temporal gyrus and sulcus, *planum temporale* (PTe), and inferior parietal regions (supramarginal gyrus), connecting to inferior frontal and more dorsal premotor regions. Connectivity to pre-supplementary and supplementary motor areas (medial regions in red and blue), basal ganglia (not shown), and cerebellum, rounds out a distributed system supporting speech. The ventral stream (blue arrow) regions primarily support lexical and semantic processing at word, sentence and narrative levels.

An extensive cortical model is also presented by Hickok and colleagues (Hickok et al., 2023). They postulate a model of speech coordination across dual processing streams which integrates primary motor cortical regions, precentral speech areas with prominent roles in speech coordination, and multiple cortical regions necessary for speech production, such as the inferior frontal gyrus (IFG) and superior temporal sulcus (STS). According to this model, speech can be construed as a distributed network across two broad processing streams: the ventral stream, which maps sound onto meaning, and the dorsal stream, which maps sound onto articulatory-based representations (Hickok & Poeppel, 2000, 2004, 2007). The left-dominant dorsal stream has a sensorimotor interface between parietal and temporal regions which projects to an articulatory network in the frontal lobe supporting articulation, specifically involving the posterior IFG (namely *pars opercularis*), the premotor cortex, and the anterior insula (Hickok & Poeppel, 2007). The bilaterally-organized ventral stream is a combinatorial lexical interface implicated in speech recognition, which connects the anterior and posterior regions of the middle temporal gyrus (MTG) and inferior temporal sulcus (ITS) (Hickok & Poeppel, 2007), extending beyond the classic left-hemisphere dominant model of speech (Tremblay & Dick, 2016). Evidence from neuroimaging studies have demonstrated an overlap in speech perception and speech production within the superior temporal gyrus (STG), incorporated in both streams (Buchsbaum et al., 2001; Hickok et al., 2003). The dual-stream model expands more simplistic models which envision speech as communication between separate regions controlling speech production and speech perception.

The contributions of the left IFG, neighboring frontal operculum, and insular cortex are emphasized by both Guenther and Hickok and colleagues. Although not addressed in Hickok and colleagues, Guenther additionally emphasizes medial frontal regions associated with motor planning, such as the pre-SMA and SMA. These regions are supported by a monosynaptic pathway known as the frontal aslant tract (Figure 2). The FAT’s connectivity of the IFG and pre-SMA/SMA, regions previously associated with motor speech properties (Hertrich et al., 2016), suggests that it may play a role in planning and initiation of speech, especially in the left hemisphere (Dick et al., 2019). Several studies have demonstrated the FAT’s association with various aspects of speech, specifically verbal fluency (Catani, Mesulam, Jakobsen, Malik, Martersteck, Wieneke, Thompson, Thiebaut de Schotten, Dell’Acqua, et al., 2013; Mandelli et al., 2014), motor speech initiation (Kinoshita et al., 2015; Zhong et al., 2022), speech fluency impairments such as stuttering (Kemerdere et al., 2016; Kronfeld-Duenias et al., 2016; Misaghi et al., 2018), and speaking rate (Jossinger et al., 2023). Few studies have explored the development of the FAT in young children. For example, Broce and colleagues (Broce et al., 2015) tracked the FAT in young children (5-8-year-olds), but found no associations with phonology and expressive language. One study has linked FAT diffusion properties to stuttering in 6-12-year-old children (Misaghi et al., 2018). However, no studies, to our knowledge, have examined developmental variability of the FAT in relation to speech in very young children, as we propose to do here.

**FIGURE 2.**
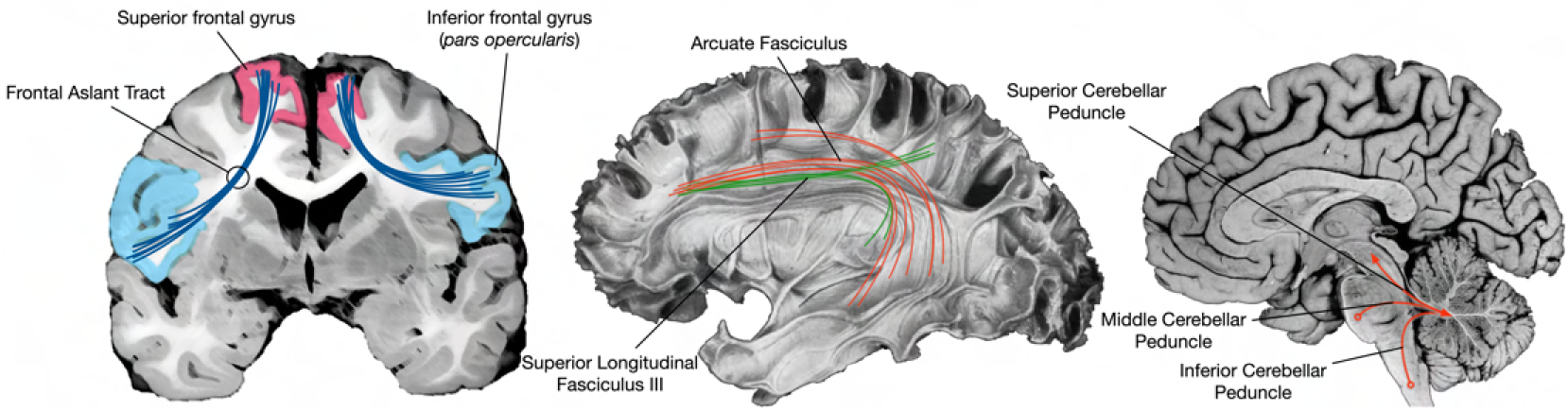
Cortical and cerebellar pathways supporting speech are examined in the current study. The frontal aslant tract (FAT) connects inferior frontal gyrus and pre-supplementary motor area; superior longitudinal fasciculus III and arcuate fasciculus connect inferior frontal and dorsal premotor cortex with supramarginal gyrus and superior and middle temporal cortex; the three brainstem peduncles support communication with cerebellum.

The left IFG is also a key node in a dorsal pathway for speech (Hickok et al., 2003; Hickok & Poeppel, 2004, 2007; Rauschecker, 1998; Rauschecker & Scott, 2009) mapping sounds onto articulatory-based representations through interactions with the posterior STG and STS (STGp and STSp), *planum temporale*, and supramarginal gyrus. The structural connectivity of these regions has been established via several invasive and non-invasive methods (Bernard et al., 2019; Willems et al., 2009). Two major fiber pathways are thought to support dorsal stream connectivity–the superior longitudinal fasciculus III (SLF III) and the arcuate fasciculus (AF). SLFIII is thought to support a fronto-parietal articulatory loop implicated in motor speech function, based on disturbances during electrostimulation (Duffau et al., 2003). SLF III is specifically defined by its connectivity with the supramarginal gyrus, involved in processing phonological inputs and outputs (Oberhuber et al., 2016), and IFG and frontal operculum (Bernal & Altman, 2009), thus establishing its putative role in speech production. AF projects more broadly, connecting posterior temporal and inferior parietal cortex with both ventral and superior frontal cortical regions (Barbeau et al., 2020). It has been proposed that the AF plays a role in speech monitoring and motor speech function, due to established connectivity with premotor and primary motor areas in the precentral gyrus (Bernal & Ardila, 2009). Hickok and colleagues have recently delineated a region possibly facilitated by this connectivity, referred to as the dorsal precentral speech area (Hickok et al., 2023). This area has been demonstrated to exert significant influence over prosodic control, an element of speech production. Supporting the claim that SLF and AF play a role in speech production is a study that demonstrated an association with AF and associated segments, including SLF III, and receptive and expressive language in children aged 5-8 years (Broce et al., 2015). The proposed study seeks to expand on previous research implicating AF and SLF III in developmental speech and language abilities by focusing on a specific component of speech, phoneme articulation, which will allow us to further parse the specific functions of these support pathways.

The cerebellum and its associated connectivity are also known to support speech and potentially speech development (Jobson et al., 2022). The involvement of the cerebellum in verbal fluency and articulation tasks is well-established (Riecker et al., 2006; Riecker et al., 2005; Schlösser et al., 1998), and anatomical evidence of functional connectivity suggests that the role of the cerebellum may go beyond simple motor generation and activation, as previously thought. As reviewed in Vias and Dick (Vias & Dick, 2017), lateral and medial regions of cerebellar grey matter have been associated with distinct aspects of speech production. Two of the cerebellar lobules that fall along the horizontal cerebellar fissure, Crus I and Crus II, have been shown to play a role in phonological processing through word generation and phonemic and verb fluency tasks (Frings et al., 2006; Schweizer et al., 2010; Stoodley et al., 2012). Activation during fMRI tasks of speech production have revealed bilateral findings in the Crus I and Crus II lobules in adults, but little is known about the development of these regions in very young children with regard to speech (Bohland & Guenther, 2006; Correia et al., 2020; Geva et al., 2021; Peeva et al., 2010; Shuster & Lemieux, 2005). The inferior, middle, and superior cerebellar peduncles (ICP, MCP, and SCP) are the three major white matter input and output pathways of the cerebellum (Naidich et al., 2009), linking the lateral cerebellar cortex with subcortical and cortical structures in the cerebral cortex (Vias & Dick, 2017). The right ICP has been linked to developmental differences in children who stutter (Johnson et al., 2022), and in adults with developmental stuttering, speech rate has been associated with white matter cellular properties of the left ICP (Jossinger et al., 2021). Additionally, white matter properties of the ICP have been correlated with articulation rate, one facet of speech production (Jossinger et al., 2021; Jossinger et al., 2023; Kronfeld-Duenias et al., 2016). In neurotypical adults, the right SCP has been associated with semantic and phonemic verbal fluency, while the right MCP is associated with speaking rate (Jossinger et al., 2023), demonstrating that these distinct cerebellar pathways contribute to elements of speech production beyond articulatory control, such as lexical access (Jossinger et al., 2023). The cerebellum has also been implicated in fine speech control during childhood for children with early speech deficits (A. Morgan et al., 2011). However, the precise anatomical structures and functional connectivity contributing to such deficits are still unknown. While the cerebellar peduncles and cerebellar grey matter have been established in aspects of speech production, further research is necessary to probe how these regions and pathways are integrated within the broader cortical speech network during development, and how this might be moderated by age.

### 1.2| Measurement of white matter properties in the developing brain

To investigate the development of neural regions and connectivity underlying speech, we examined changes in the microstructural properties of neural and glial cells, as well as their processes (e.g., axons and dendrites). These changes influence the restricted and hindered components of the diffusion signal, which are shaped by cellular properties such as size, density, orientation, and myelination. By leveraging restriction spectrum imaging (RSI) reconstruction, we can separate restricted diffusion from hindered diffusion and free water diffusion (Brunsing et al., 2017; Palmer et al., 2022; White, Leergaard, et al., 2013a; White, McDonald, Farid, et al., 2013; White et al., 2014).

The restricted diffusion signal reflects water diffusion confined within cell bodies and processes, providing insights into *intracellular* diffusion properties. In contrast, hindered diffusion captures the characteristics of water navigating the extracellular environment, which constrains molecules to follow more winding or “tortuous” paths in both grey and white matter. Modeling the hindered diffusion signal allows us to better characterize these extracellular properties. To achieve this level of analysis, we used a specialized multi-shell high-angular-resolution diffusion imaging (HARDI) protocol. This protocol includes both low (b = 500 s/mm^2^) and high (b = 3000 s/mm^2^) b-values, robustly sampling the diffusion signal at high b-values. High b-values are especially sensitive to diffusion properties that evolve over longer time intervals. For example, the average diffusion coefficient (ADC) of restricted intracellular water decreases with diffusion time, but this sensitivity diminishes at lower b-values, where overlap with hindered diffusion becomes more prominent. This combination of advanced acquisition and RSI reconstruction enables improved separation of hindered and restricted diffusion compared to traditional approaches, such as diffusion tensor imaging (DTI) or single-shell HARDI. It also offers a performance comparable to other multicomponent models, such as Neurite Orientation Density and Dispersion Imaging (NODDI).

This methodological foundation enables the indirect measurement of “cellularity,” a term that broadly encompasses the microstructural properties influencing restricted and hindered diffusion signals. While the diffusion signal correlates with these microstructural changes, it is important to note that these properties cannot be directly observed. The restricted diffusion component is expected to increase with processes such as enhanced myelination in both white and grey matter (which reduces extracellular space volume and axonal membrane permeability), increased neurite diameter, dendritic sprouting, or the recruitment and activation of microglia. In contrast, the hindered diffusion component is likely to decrease under the same conditions, reflecting the extracellular environment.

Although these observations are indirect inferences drawn from the diffusion signal, prior research demonstrates that such changes occur in both white and grey matter and can be detected using the proposed diffusion protocol and RSI reconstruction (Palmer et al., 2022). These findings are further supported by histological evidence (White, Leergaard, et al., 2013a).

Traditionally, DWI studies have focused on understanding white matter development in specific fiber pathways, or in white matter defined broadly (e.g., via analysis of tract-based spatial statistics). These approaches generally focus on measuring anisotropic diffusion within axons, typically using diffusion-tensor or higher-order spherical deconvolution models. For example, many studies conducting tractography examine fractional anisotropy (FA) differences across age (Lebel et al., 2017), or in association with individual differences in performance on some task (Catani, Mesulam, Jakobsen, Malik, Martersteck, Wieneke, Thompson, Thiebaut de Schotten, Dell’Acqua, et al., 2013; Dick et al., 2019). An advantage of RSI is that we can measure both white matter tracts and grey matter of the cortex, subcortical structures, and cerebellum (Palmer et al., 2022). While other measures like mean diffusivity are sensitive to differences in grey matter (Sagi et al., 2012), RSI provides a more detailed and biologically specific characterization of tissue properties by distinguishing intra- and extra-cellular diffusion components. The normalized total signal fractions for restricted (RNT) and hindered (HNT) diffusion produced by this method can be interpreted as reflecting the relative contributions of maturational and developmental cellular processes associated with cell bodies and neurites to diffusion signal in the different compartments. Thus, we can measure the relationship between RSI metrics and cellularity in both grey and white matter structures of the speech network that can be related to individual behavioral differences.

With RSI, we can examine individual differences in both grey and white matter. However, we can also apply more sophisticated analysis to the white matter tracts themselves. We do this in the present study with Automated Fiber Quantification (AFQ; (Yeatman et al., 2012)). Unlike traditional tractography methods that average across the entire white matter bundle, AFQ segments each tract into 100 equidistant nodes, allowing us to establish a “tract profile” for each child and observe individual differences at specific positions along the tracts (Yeatman et al., 2012), and how these individual differences are associated with behavior. For the current analysis, we are interested in white matter pathways associated with the neural speech network, specifically the FAT, AF, SLF III, and superior, middle, and inferior cerebellar peduncles (SCP, MCP, ICP). It is anticipated that these pathways will display differences in measured restricted and hindered diffusion values along the white matter tracts based on speech performance measured by a phoneme articulation task.

### 1.3| Variability of speech production in developmental samples, including those at risk for speech delay or disorder

In the present study, we are interested in examining individual differences in speech production, and their association with structural brain development as measured by diffusion imaging. Examining this relationship requires measured variability in both microstructural properties of the brain, and performance on a behavioral task. In order to expand the degree of variability in speech performance, we included as part of the typically developing (TD) sample a sample of children diagnosed with attention-deficit/hyperactivity disorder (ADHD). The reason for the inclusion of this sample of children was two-fold. First, it was a practical choice given our data collection of the TD sample was part of a broader study that included children with ADHD. The first goal is thus to substantially increase the sample size and statistical power. However, the second goal is to increase the variability in performance on the outcome measure of speech production.

The literature provides evidence that children with ADHD are are more likely to display speech production errors than their typically developing peers. For example, children with ADHD are at increased risk for comorbid speech delay (McGrath et al., 2008) and other speech and language impairments (Baker & Cantwell, 1992; Blood et al., 2003; Damico et al., 2010; Donaher & Richels, 2012; Druker et al., 2019; Healey & Reid, 2003), such as specific language impairment. Mueller and Tomblin (Mueller & Tomblin, 2012; Tomblin & Mueller, 2012) reviewed more than 20 studies that evaluated the comorbidity of ADHD and speech and language impairment throughout childhood, finding on average across studies that 50% of children with ADHD are also diagnosed with speech and / or language disorders. In fact, speech, language, and communication difficulties are among the most common comorbid diagnoses for children with ADHD (Mueller & Tomblin, 2012). Specifically for our purposes, children with ADHD symptoms are more likely to display speech *production* difficulties compared to their typically developing peers (Lee et al., 2017), and they perform more poorly than typically developing children on articulation and phonology tasks (Sariyer et al., 2023). Thus, we expect that the ADHD group will contribute to the variability in our measure of interest, in addition to increasing the sample size. This sample makeup offers a distinct and valuable background for delineating the developmental trajectories of various brain regions and fiber pathways involved in speech.

### 1.4| The present study

The primary aim of the present study is to probe how the neural systems that implement speech develop structurally in children with a wide variety of speech production abilities, including children at risk of speech disorders, such as those with ADHD. A validated measure of phoneme articulation known as the Syllable Repetition Task (SRT) was used to explore the specific role of these regions of interest (ROIs). The SRT is a speech production task suitable for young children with limited phonemic inventories (Shriberg & Lohmeier, 2008; Shriberg et al., 2009), making it an appropriate task to measure speech production errors in young children with and without ADHD. It reliably measures expressive language impairment and auditory-perceptual speech processing errors (Shriberg et al., 2009). The task evaluates pre-articulatory planning, phonological planning, and transformation of phonological plans into motor speech execution (Levelt et al., 1999; Rvachew & Matthews, 2017).

We predict that 1) performance on the SRT will be associated with grey matter cellularity in the cortical and subcortical regions identified to be involved in the neural network of speech; 2) these associations will be moderated by age; and 3) performance on the SRT will be associated with white matter microstructural properties in the fiber pathways involved with speech speech production, including the FAT, AF, SLF III, and the three cerebellar peduncles.

## 2| METHOD

### 2.1| Participants

This study is a substudy of a broader research project examining 322 4-7-year-olds. That study involved MRI scanning and collection of various behavioral and clinical measures. Roughly half of that broader study sample had a diagnosis of ADHD. For this substudy, 113 children also completed the SRT outside the MRI scanner (the COVID pandemic prohibited continued data collection of this particular measure on the full sample because the head-mounted microphone was a transmission risk). We focus in this substudy on the sample of 113 participants who completed the task.

Of the 113 participants who completed the SRT, 10 participants were excluded based on excessive movement (see Image Acquisition below), and 9 participants were excluded due to being left-handed (defined by the Edinburgh Handedness Inventory) (Oldfield, 1971). Thus, the final analyzed sample size was *n* = 94 (*M_age_* = 5.5 years, *SD* = 0.82, 70 males).

#### Recruitment and Eligibility Requirements

The study took place in a large urban southeastern city in the U.S. with a large Hispanic/Latino population. Children and their caregivers were recruited from local schools, open houses/parent workshops, mental health agencies, and radio and newspaper ads. Exclusionary criteria for the children included intellectual disability (IQ lower than 70 on the Wechsler Preschool and Primary Scale of Intelligence 4th edition; WPPSI-IV), confirmed history of Autism Spectrum Disorder, and currently or previously taking psychotropic medication, including children who have been medicated for ADHD. The study was reviewed an approved by the Florida International University Institutional Review Board.

#### Demographics of the Sample

Within this sub-sample, there were 47 typically-developing children and 47 children who were diagnosed with ADHD. ADHD diagnosis was accomplished through a combination of parent structured interview (C-DISC (Shaffer et al., 2000) and parent and teacher ratings of symptoms and impairment (Disruptive Behavior Disorders Rating Scale (DBD), which involves teacher and parent reporting of symptoms (Fabiano et al., 2006a; Graziano et al., 2022), and Impairment Rating Scales (Fabiano et al., 2006b). The DBD, updated for DSM-5 terminology, assesses for symptoms of ADHD, including hyperactivity and impulsivity (Graziano et al., 2022). Academic, behavioral, and social impairments were measured by a score of three or higher on a seven point Impairment Rating Scale (Graziano et al., 2022). For this study, ADHD was not considered categorically in our statistical models, but we controlled for ADHD symptoms of inattention and hyperactivity, as measured by the DBD.

The demographic breakdown of the subsample, which is based on United States National Institutes of Health demographic categories, was: 82% Hispanic/Latino; 87% White; 7% Black; 3% Asian; 3% More than one race. The distribution of maternal education in AHEAD also suggests a good range in our sample catchment: 8.6% of mothers had a high school degree or less, 15.2% had some college, 13.0% had associate’s degrees, 25.0% had bachelor’s degrees and 38.0% had an advanced degree.

### 2.2| Experimental Paradigm

Children completed an MRI scan and the SRT on the same day, as described below.

#### Syllable Repetition Task (SRT)

The SRT is a child-friendly speech production task that examines expressive speech abilities in individuals with limited phonetic inventories, such as young speakers (Shriberg & Lohmeier, 2008). Research suggests that the SRT is an accurate and stable task for measuring expressive language impairment, as well as auditory-perceptual speech processing errors (Shriberg et al., 2009). The task measures pre-articulatory planning and subsequent planning of articulatory gestures prior to and including motor execution of speech, which includes pre-articulatory encoding and memory of speech sounds, phonological planning, and transforming the phonological plan into a motor plan (Levelt et al., 1999; Rvachew & Matthews, 2017).

During the task, children were asked to repeat a list of 18 two-to four-syllable nonsense words which were played over computer speakers. The task is available as part of a power-point presentation with the audio embedded in each slide, which includes text. The experimenter ensured that the slide presentation was turned away from the participant so that children who could read did not use orthographic information to complete the task. The task starts with simple, two-syllable nonwords such as “ba-da”, and increases in difficulty to nonwords such as “ba-na-ma-da”. Children were recorded during the task with a head-mounted microphone.

The recordings were independently transcribed and scored by three Spanish-English bilingual research assistants and scored according to the SRT scoring manual, which provided a final percentage score for proportion of correct syllables spoken. Syllable additions, or syllables that the participant produced that were not part of the target word, were tallied. If four or more responses included syllable additions (25% of items), the score was deemed invalid. In addition, given the large proportion of bilingual Spanish/English speaking children enrolled in our study and since the SRT was developed for monolingual English-speaking participants, we decided to score the SRT using both the traditional scoring manual as well as a modified bilingual scoring system. The bilingual scoring method modified the traditional scoring method to allow for b/v substitutions, which are two separate phonemes in English but are commonly both pronounced as “b” in Spanish.

Three transcribers, all Spanish-English bilinguals, were trained by a licensed Spanish-English bilingual speech-language pathologist (author A.R.A.). The three transcribers scored according to the bilingual scoring method and had a moderate inter-rater reliability score of *κ* = 0.587 (Cohen, 1960). In cases of disagreement, the score for each word was determined by majority (2 out of 3) vote. If agreement could not be reached, the recording was replayed with all transcribers and a licensed Spanish-English bilingual speech-language pathologist present for a final determination. The bilingual scoring was used in the present analysis.

The sample provided a good range of scores on the SRT, providing sufficient variability to examine associations with our brain measures. Performance on the percent correct syllables ranged from 22 to 100, with a mean percentage of 83.67% (*SD* = 14.14; see Figure 3). There was an overall positive effect of age (Figure 3a). As expected, older children performed better on the SRT (*r* = 0.25, *p* = .015). We also examined whether there were performance differences between children diagnosed with ADHD and typically developing children. There was no significant performance difference between the groups *t*(92) = −1.20, *p* = 0.23, so we did not examine the group in the analysis. Instead, as described in the analysis section, we controlled for inattention and hyperactivity symptoms. However, the density plot (Figure 3b) revealed that children with ADHD did contribute scores on the lower end of the performance range, as predicted by literature suggesting children with ADHD display speech production errors. Thus, the inclusion of the ADHD sample did contribute to variability on the measure, and improved the sample size by 100%.

**FIGURE 3.**
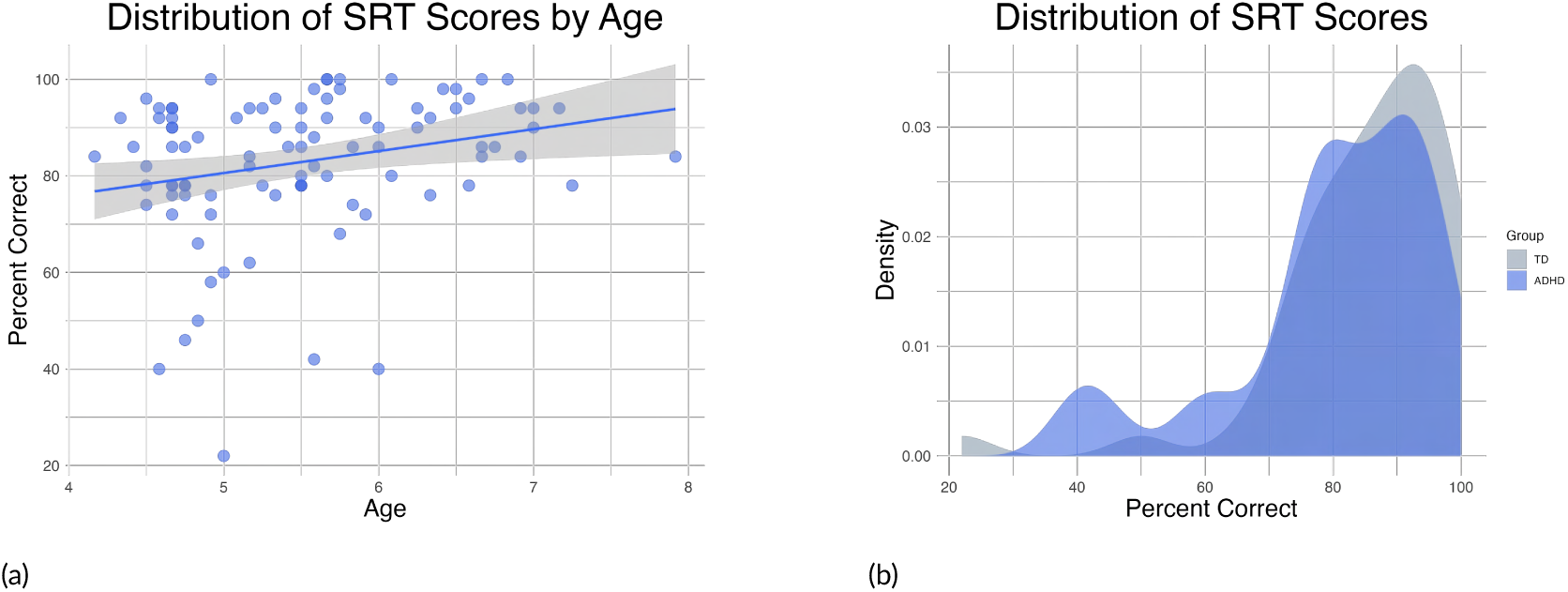
Distribution of Syllable Repetition Task (SRT) scores. (A) Performance by age group. (B) Density plot showing the overall distribution of scores. Both plots display scores as percentage correct using the bilingual scoring system, revealing a good range of performance with some degree of negative skew. Figure 3a shows a positive effect of age, demonstrating that older children performed better on average. Figure 3b reveals that while there was no significant difference between groups, children with ADHD did contribute scores on the lower end of the performance spectrum.

### 2.3| Image Acquisition

All imaging was performed using a research-dedicated 3-T Siemens MAGNETOM Prisma MRI scanner (V11C) with a 32-channel coil located on the university campus. Children first completed a preparatory phase using realistic mock scanner in the room across the hall from the magnet. Here, they were trained to stay still and were also acclimated to the enclosed space of the magnet, to the back projection visual presentation system and to the scanner noises (in this case, presented with headphones). When they were properly trained and acclimated, they were moved to the magnet. In the magnet, during the structural scans, children watched a child-friendly movie of their choice. Sound was presented through MRI-compatible headphones that also functioned as additional ear protection.

T1-weighted and DWI scans were acquired for each participant. T1-weighted MRI scans were collected using a 3D T1-weighted inversion prepared RF-spoiled gradient echo sequence (axial; TR/TE 2500/2.88; 1 x 1 x 1 mm, 7 min 12 s acquisition time) with prospective motion correction (Siemens vNav; (Tisdall et al., 2012)), according to the Adolescent Brain Cognitive Development (ABCD) protocol (Hagler Jr et al., 2019).

Movement artifacts pose challenges for T1-weighted images, which affects registration in the diffusion scan. Each T1 image was thoroughly reviewed by A.S.D. We applied a visual rating system ranging from ‘poor = 1’ to ‘excellent = 4’ for each T1-weighted image, with half-point allowances (e.g., 3.5). Most images were in the 3-4 range, with an average image rating of 3.63 (*SD* = 0.53)

DWI scans were acquired via high-angular-resolution diffusion imaging (HARDI) with multiband acceleration factor 3 (EPI acquisition; TR/TE = 4100/88 ms; 1.7 x 1.7 x 1.7 mm; 81 slices no gap; 96 diffusion directions plus 6 b = 0 s/mm^2^: b = 0 s/mm^2^ [6 dirs], b = 500 s/mm^2^ [6-dirs], 1000 s/mm^2^ [15-dirs], 2000 s/mm^2^ [15-dirs] and 3000 s/mm^2^ [60-dirs] s/mm^2^; A-to-P direction [7 m 31 s acquisition time]). A second brief scan at b = 0 s/mm^2^ in P-to-A direction was acquired to help deal with susceptibility artifacts.

#### Image Post-Processing

DWI preprocessing was performed using a number of complementary software suites. The steps were as follows: 1) outlier detection and replacement, and volume-to-slice correction using a synthetized DWI model, implemented in FSL eddy (Andersson et al., 2017; Andersson et al., 2016) (note motion and eddy current distortion correction are implemented in the next step); 2) motion and eddy-current distortion correction, implemented with TORTOISE DIFFPREP (Barnett et al., 2014) instead of FSL eddy; 3) creation of a synthesized T2-weighted image from the T1-weighted scan using Synb0-DisCo (Schilling et al., 2020; Schilling et al., 2019); 4) correction of spatial and intensity distortions caused by B_0_ field inhomogeneity, using TORTOISE DRBUDDI (Irfanoglu et al., 2015) implementing blip-up/blip-down distortion correction (Andersson et al., 2003; H. Chang & Fitzpatrick, 1992; D. Holland et al., 2010; P. S. Morgan et al., 2004). This step uses both the reverse-phase encoded b = 0 s/mm^2^ image for the estimation of the field map, and the synthesized T2-weighted image for imposition of geometry constraints; 5) gradient non-linearity correction (gradwarp) using the gradient coefficients supplied by Siemens (Bammer et al., 2003; Barnett et al., 2021; Glover & Pelc, 2019). The outlier-replaced, slice-to-volume registered, transformed for motion, eddy-current corrected, b0-induced susceptibility field corrected, gradient non-linearity corrected images are resampled to the T1-weighted resolution (to 1 mm^3^) and registered to the T1-weighted image in a single interpolation step. For whole-brain analysis, data are warped to the ABCD atlas space, and reported in LPS atlas coordinates (L = +; P = +; S = +), which are derived from the DICOM coordinate system.

### 2.4| Diffusion Metrics

#### Restriction Spectrum Imaging

We took advantage of the multi-shell HARDI acquisition to implement a reconstruction of the diffusion signal with the RSI model (Brunsing et al., 2017; White, Leergaard, et al., 2013b; White, McDonald, Farid, et al., 2013; White et al., 2014). RSI reconstruction was accomplished in MATLAB using the model from White and colleagues (Brunsing et al., 2017; White, Leergaard, et al., 2013b; White, McDonald, Farid, et al., 2013; White et al., 2014), and Hagler and colleagues (Hagler Jr et al., 2019), which was updated in Palmer and colleagues (Palmer et al., 2022). The RSI model can be used to quantify the relative proportion of restricted, hindered, and free water diffusion within each voxel of the brain. These components–restricted, hindered, and free water–have intrinsic diffusion characteristics. Free water (e.g., cerebrospinal fluid) is water diffusion unimpeded by tissue structure. In biological tissue, though, the two additional modes of hindered and restricted diffusion predominate (Bihan, 1995). Hindered diffusion describes the behavior of water hindered by the presences of neurites, glia, and other cells, and follows a Guassian displacement pattern. The signal originates from both extracellular and intracellular spaces with dimensions larger than the diffusion length scale (10 *µ* m), and it is affected by the length of the diffusion path by which molecules must travel to navigate cell obstructions. On the other hand, the primary source of restricted diffusion signal in biological tissue comes from cell membranes. Restricted diffusion thus relates to the physical obstruction of molecules within cellular compartments. If the diffusion time Δ is long enough, the length scale of diffusion will vary depending on whether diffusion is hindered or restricted (White, Leergaard, et al., 2013b). The acquisition parameters applied in the present study are designed to optimize sensitivity to the different diffusion processes.

A number of quantitative metrics can be recovered from the RSI reconstruction of the diffusion data (Palmer et al., 2022). The model estimates diffusion within different compartments, and provides metrics that are normalized into signal fractions in order to determine the relative proportion of restricted, hindered, and free water diffusion within each voxel. We focus on two metrics–the restricted normalized total signal fraction (RNT) and the hindered normalized total signal fraction (HNT).

The hindered and restricted compartments are modeled as fourth-order spherical harmonic functions and the free water compartment is modeled using zeroth-order spherical harmonic functions. Axial diffusivity is assumed to be constant at 1 x 10^−3^ mm^2^/s for the restricted and hindered compartments. For the restricted compartment, the radial diffusivity is fixed to 0 mm^2^/s. For the hindered compartment, radial diffusivity is fixed to 0.9 x 10^−3^ mm^2^/s. Isotropic diffusivity is fixed to 3 x 10^−3^ mm^2^/s for the free water compartment. Spherical deconvolution is applied to reconstruct the fiber orientation distribution (FOD) in each voxel for the restricted compartment. The norm of the second and fourth order spherical harmonic coefficients is the restricted directional measure (RND; modeling oriented diffusion from multiple directions in a voxel), and the spherical mean of the FOD across all orientations is the restricted isotropic measure (RNI). The sum of RND and RNI is the RNT. For more details, see (Brunsing et al., 2017; Palmer et al., 2022; White, Leergaard, et al., 2013b; White, McDonald, Farid, et al., 2013; White et al., 2014).

The two metrics extracted from RSI, RNT and HNT, can be interpreted as reflecting the relative contributions of maturational and developmental cellular processes associated with cell bodies and neurites (neural projections from the cell body such as axons and dendrites) to signal in the different compartments. These maturational and developmental processes include myelination, dendritic sprouting, changes in neurite diameter with constant neurite density, changes in cell body size with constant cell density, and changes in the concentration of mature astrocytes. Inspection of the diffusion maps of these metrics, and prior research using the same acquisition parameters in adolescents (Palmer et al., 2022), show that RNT and HNT reliably distinguish white and grey matter and show age-related change in both tissue types (Palmer et al., 2022). The two metrics tend to be anti-correlated. For example, increased myelination is associated with increased restricted diffusion (due to reduced permeability of axonal membranes), and decreased hindered diffusion (due to reduced volume of extracellular space). However, this is not always the case, and depends on specific tissue properties and the parcellation of signal within the different diffusion compartments in a voxel.

## 3| DATA ANALYSIS

### Statistical Model for FEMA and AFQ Analysis

We conducted the statistical analysis using both a fast and efficient mixed-effects algorithm, FEMA(Parekh et al., 2024) and Automated Fiber Quantification, AFQ(Yeatman et al., 2012). FEMA is a voxelwise wholebrain analysis method, and AFQ is designed for specific fiber pathways. These methods are described below. First we detail the statistical models.

The same basic model was run for three analyses: 1) the FEMA Model Main effect tested, voxelwise, the association between SRT performance and diffusion metrics; 2) the FEMA Model Interaction added the interaction term for age and SRT performance, which tests, voxelwise, where the association between SRT and diffusion metrics are moderated by age; 3) the AFQ model, which conducts the AFQ analysis to identify *where* along the specific fiber pathway performance differences are associated with diffusion metrics (by entering the interaction of SRT performance and NodeID). These models are described in Equations 1-6, and were run with RNT and HNT as outcomes, and the specified covariates. Each method is further detailed below.

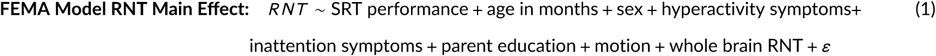

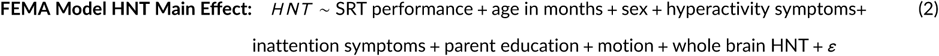

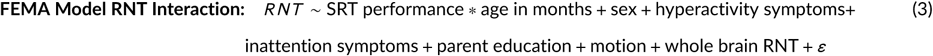

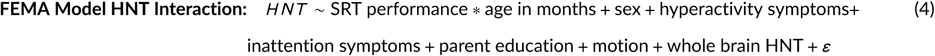

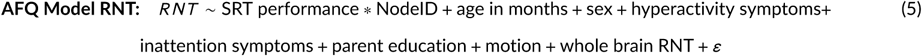

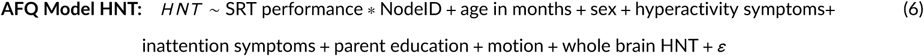

#### Covariates

We included age, sex assigned at birth (sex), parental education, and ADHD symptomology (inattention and hyperactivity from the DBD (Fabiano et al., 2006a; Graziano et al., 2022)). Age, sex, and parental education were all parent reported, and the DBD measures also included teacher report. In addition, we included the whole brain diffusion metric (either RNT or HNT). Whole-brain diffusion data were output directly from the post-processing stream to control for general individual differences in diffusion properties of the brain.

Finally, we know participant movement has a substantial effect on DWI measurements, and can introduce spurious group differences in cases where none are present in the biological tissue (Yendiki et al., 2014). Therefore, incorporating movement as a nuisance regressor is recommended (Yendiki et al., 2014). To do this, we estimated movement using the root mean square (RMS) output from FSL eddy and implemented an overall movement cutoff for inclusion in analysis. This was arbitrarily defined as average RMS movement of more than one voxel (1.7 mm) over the course of the scan. As noted above, nineteen children (6% of the overall sample of *n* = 322) exceeded the movement criteria and were dropped from the sample. For participants retained in the analysis, movement was incorporated as a nuisance regressor in all analyses.

#### Fast and Efficient Mixed-Effects Algorithm (FEMA)

We performed a whole brain, voxelwise analysis using FEMA. The algorithm is uniquely suited for large sample sizes due to its ability to efficiently perform whole-brain image-wise analyses on complex large scale imaging data (Parekh et al., 2024). It was chosen as an appropriate analysis for the current study because it is optimized for the ABCD brain atlas, and the T1-weighted MRI scans for the current study were collected according to protocol established by the ABCD study (Hagler Jr et al., 2019). The design matrix was constructed to examine the effect of SRT performance on our diffusion metrics of interest, as specified in the statistical models above. The per-voxel threshold was set at *p* < 0.005. For the interaction effect, the same basic results were obtained at *p* < 0.001, so this more stringent threshold is reported for that analysis. In order to apply cluster-mass correction to correct for multiple comparisons, we repeated the analysis using FSL’s randomise (note since this is not a multilevel model, the same results are obtained in FSL and FEMA). Both uncorrected and cluster-mass-corrected data are reported.

#### Automated Fiber Quantification (AFQ)

Our second analysis allowed us to examine our white matter tracts of interest in-depth via Automated Fiber Quantification (AFQ). The pre-processed DWI images were analyzed by the AFQ software developed by Yeatman and colleagues (Yeatman et al., 2012), which has since been applied in a number of studies of speech fluency (Jossinger et al., 2021; Jossinger et al., 2023; Kronfeld-Duenias et al., 2016; Yablonski et al., 2021). Using the AFQ toolbox, we identified and quantified 6 bilateral white matter tracts for analysis: FAT, AF, SLFIII, ICP, MCP, and SCP. Parameters followed the AFQ protocol developed by Kruper and colleagues (Kruper et al., 2021), with the addition of the bilateral frontal aslant tract as described by Kronfeld-Duenias and colleagues (Kronfeld-Duenias et al., 2016).

The implementation of the AFQ software, as described in Yeatman and colleagues (Yeatman et al., 2012), begins with the identification of regions of interest. Fiber tracts are identified using a probabilistic streamline tracking algorithm and refined based on waypoint regions of interest, which define the trajectory of the fascicle. An iterative procedure cleans the bundles by removing fibers that are more than 4 standard deviations above the mean fiber length or 5 standard deviations from the core of the fiber tract. AFQ software calculates the tract profiles with a vector of one hundred values representing diffusion properties, which have been sampled at equidistant locations along the central portion of the tract.

Automatic segmentation of the FAT and SLF III was carried out using the standard AFQ regions (Yeatman et al., 2012). On visual inspection, we established new seed ROIs for the AF based on the long segment definition, as defined in (Broce et al., 2015). For the cerebellar peduncles, we used the revised protocol from Jossinger and colleagues (Jossinger et al., 2023). New ROIs defined on the Montreal Neurological Institute (MNI) template were used to identify the peduncles, together with previous ROIs established by Bruckert and colleagues (Bruckert et al., 2019). This method uses probabalistic tractography coupled with CSD modeling, which has been shown to be more successful in tracking the decussation of the cerebellar peduncles, compared to the traditional deterministic tractography such as that used by Bruckert and colleagues (Bruckert et al., 2019; Jossinger et al., 2023). Upon visual inspection of the MCP, we modified the regions of interest to remain consistent with established definitions (Jobson et al., 2022; Nagahama et al., 2021).

After running the AFQ analysis, all data were retained for left and right FAT, SLF III, and AF. All participants were retained for left ICP, but for right ICP, 1 participant did not have a complete tract profile. For the left MCP, 3 participants had incomplete tract profiles, and 1 participant had an incomplete tract profile for the right MCP. The SCP proved to be the most difficult to track, as 39 participants had incomplete tract profiles (41 %) for the left SCP, and 32 participants had incomplete tract profiles (34 %) for the right SCP. For all incidents of missingness, we performed deletion of the individual tract profile for that participant (note that these participants were retained in the whole brain analysis).

From these individual profiles, we created standardized tract profiles by calculating the mean and standard deviation of each diffusion property at each node of each tract, which we then applied to a series of generalized additive models (GAMs). GAMs have been shown to be well-suited for modeling diffusion data based on AFQ, since they account for the distribution and non-continuous nature of diffusion metrics (Muncy et al., 2022). Using this method, we identified nodes which significantly differed in diffusion properties along each white matter tract, and related these differences to SRT performance. We compared two different models to determine the best fit: one simple model calculating the main effect of SRT score on the diffusion metrics, and one that took into account the interaction of the nodes, to determine the best model fit for the current analysis. For graphing purposes, we created groups based on a median split. This allowed us to plot how diffusion metrics differed between high and low SRT scores at different node intervals. In the statistical models, SRT was entered as a continuous variable.

## 4| RESULTS

### 4.1| Whole Brain Analysis

We first present results for the four voxelwise FEMA models. As a brief summary, for all models only the cerebellar findings survive correction for multiple comparisons. We report uncorrected findings because several clusters were found in expected regions based on our literature review, so we focus on these clusters. In the Discussion, we consider these findings with this caveat in mind. Regions and pathways are from the ABCD atlas space, using the Destrieux Freesurfer regions (Destrieux et al., 2010) based on the anatomical definitions from Duvernoy (Naidich et al., 2009), and the pathways from Hagler et al. (Hagler et al., 2009).

Figures 4 and 5 demonstrate the main effect, reporting voxels that show an association between SRT performance and either RNT or HNT. For RNT, uncorrected (Figure 4 A) whole brain voxelwise analysis revealed that speech performance on the SRT task was associated with grey and white matter cellularity in multiple regions, including bilateral IFG *pars opercularis*, right pre-SMA/SMA, FAT white matter, and bilateral cerebellum grey and white matter (the cerebellar peduncles). As Figure 4 B shows, only the cerebellar clusters survived the cluster correction. For HNT, the same regions were revealed to be associated with SRT performance. However, the cerebellar clusters were smaller, and in fact no clusters survived statistical correction (Figure 4 A and B).

**FIGURE 4.**
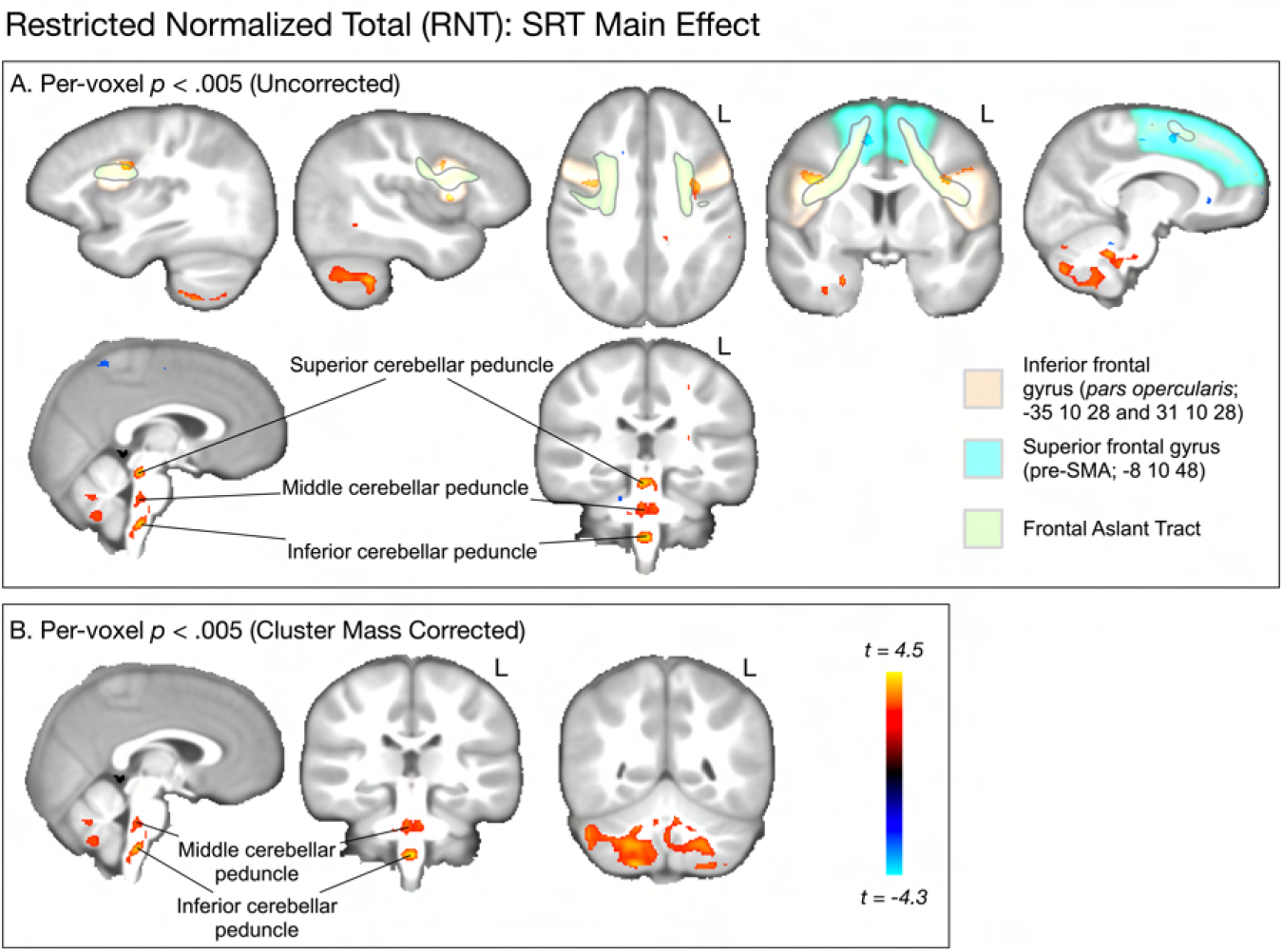
Whole brain, voxelwise analysis of the main effect association between SRT performance and RNT (Model Equation 1). A. Uncorrected (*p* < .005) clusters are highlighted for regions and pathways reviewed in the introduction. Areas in red spectrum indicate a positive association between SRT performance and grey and white matter cellular properties, as measured by RNT. Blue indicates a negative association between SRT performance and RNT. B. Cluster mass corrected (*p* < .005) results show only clusters in cerebellar grey and white matter.

**FIGURE 5.**
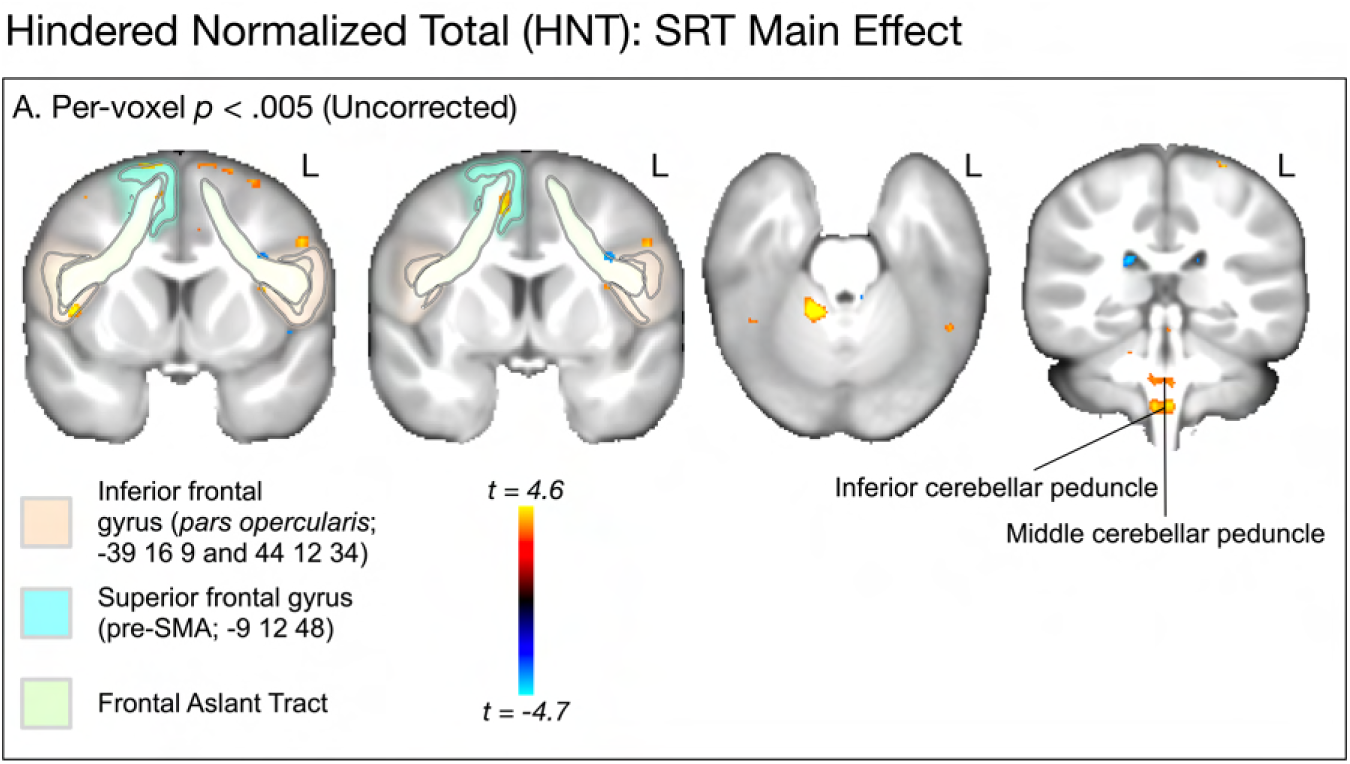
Whole brain, voxelwise analysis of the main effect association between SRT performance and HNT (Model Equation 2). A. Uncorrected (*p* < .005) clusters are highlighted for regions and pathways reviewed in the introduction. Areas in red spectrum indicate a positive association between SRT performance and grey and white matter cellular properties, as measured by RNT. Blue indicates a negative association between SRT performance and RNT. No voxels in the whole brain analysis survived cluster mass correction (*p* < .005).

Figures 6 and 7 show the results for the interaction effect. Note these are reported for the more strict *p* < .001 threshold, as the cluster-corrected findings were the same for the *p* < .005 level. Age moderated the association between SRT and both RNT and HNT in left frontal aslant tract, bilateral caudate, right globus pallidus, and large parts of bilateral cerebellum. For RNT, there was an additional cluster in left *pars opercularis*. In both RNT and HNT analyses, only the cerebellar clusters survived multiple comparison correction (B in both Figures). Figure 8 shows the nature of this interaction for the clusters revealed in cerebellum. For RNT, the interaction effect shows that performance is positively associated with RNT, especially for older ages. For HNT, the performance association reduces as age increases.

**FIGURE 6.**
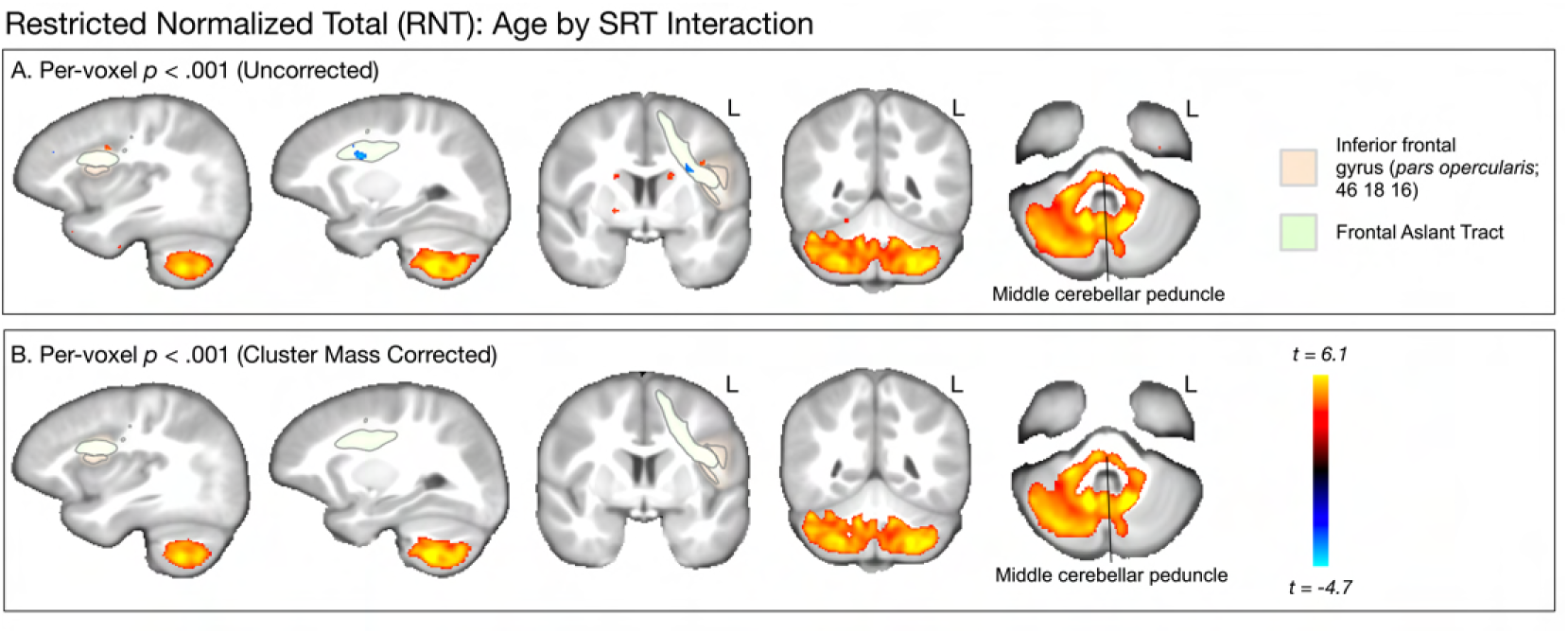
Whole brain, voxelwise analysis of the interaction effect showing how age moderates the association between SRT performance and RNT (Model Equation 3). A. Uncorrected (*p* < .001) clusters are highlighted for regions and pathways reviewed in the introduction. Areas in red spectrum indicate a positive interaction slope, and blue indicates a negative interaction slope. B. Cluster mass corrected (*p* < .001) results show only clusters in cerebellar grey and white matter.

**FIGURE 7.**
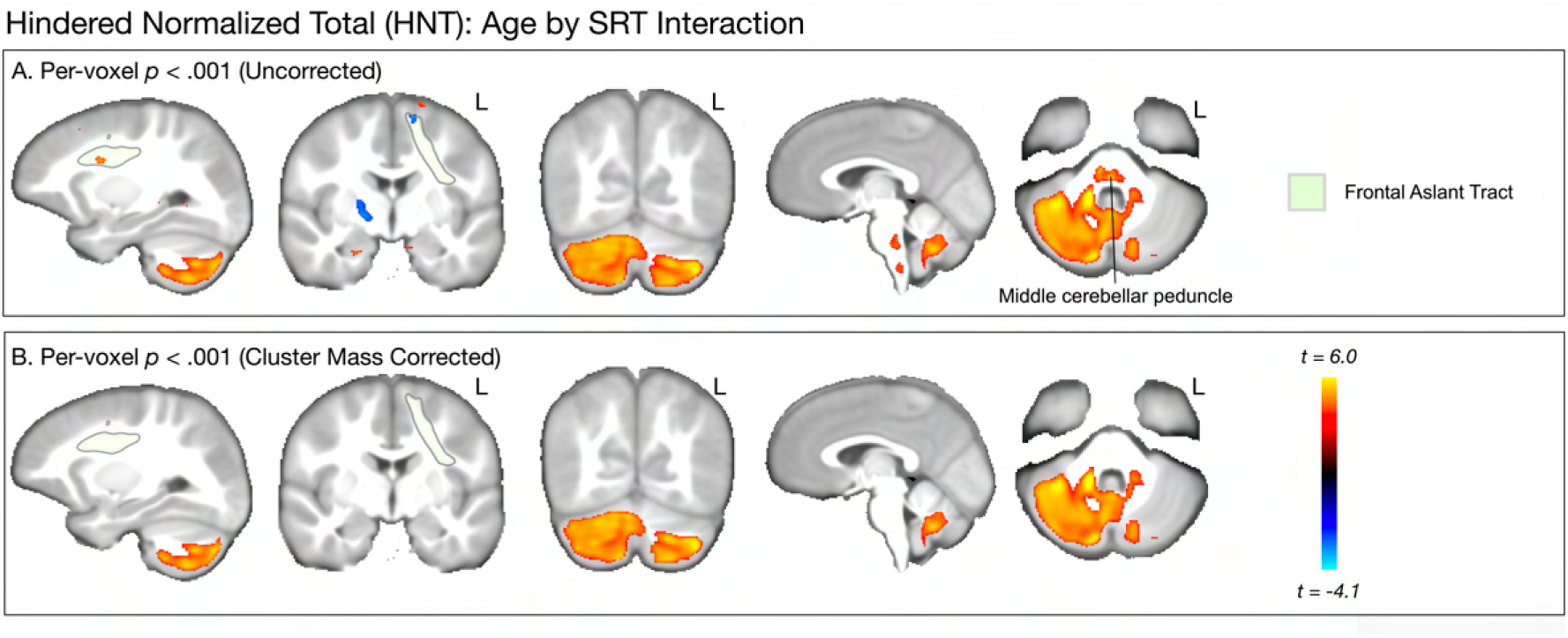
Whole brain, voxelwise analysis of the interaction effect showing how age moderates the association between SRT performance and HNT (Model Equation 4). A. Uncorrected (*p* < .001) clusters are highlighted for regions and pathways reviewed in the introduction. Areas in red spectrum indicate a positive interaction slope, and blue indicates a negative interaction slope. B. Cluster mass corrected (*p* < .001) results show only clusters in cerebellar grey and white matter.

**FIGURE 8.**
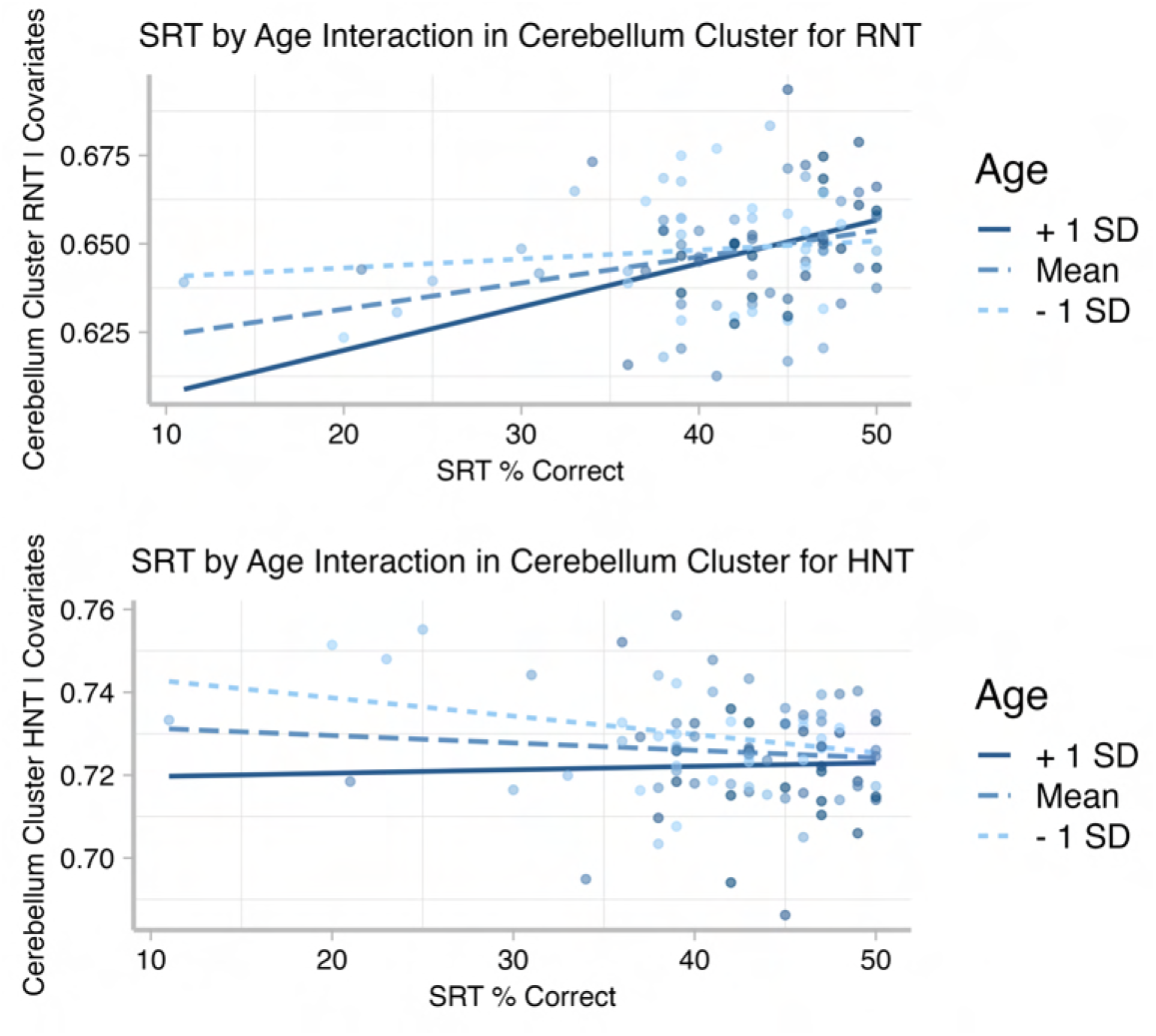
Residualized scatterplots show that age moderates the association between (top) SRT performance and Restricted Diffusion; and (bottom) SRT performance and Hindered Diffusion in the cerebellum. Data are recovered from the cluster-corrected voxels for the whole-brain interaction analysis, for each participant.

### 4.2| AFQ Analysis

To further probe the relationship between SRT performance and diffusion metrics along each white matter tract, we examined the tract profiles of white matter pathways identified in our literature review as potentially important for speech. Thus, AFQ analysis was conducted on the FAT, SLFIII, AF, ICP, MCP, and SCP.

Our primary aim for this analysis was to examine the association between SRT performance and diffusion properties. Since there are 100 nodes along the tract profile, rather than conducting separate tests for each node, we examined each tract in a series of GAM models. We were mainly interested in whether SRT performance was moderated by placement (NodeID) along the tract profile. That is, we sought to examine whether performance differences were evident at certain points along the tract, but not others. This is the interaction effect.

Unlike traditional linear models, GAMs do not return a single, unified statistic summarizing the interaction. Instead, the interaction can be assessed through changes in the model fit (e.g., via deviance explained or likelihood ratio tests) and visually interpreted through plots of the fitted curves. Statistical significance can also be evaluated at specific points (knots) along the trajectory. Thus, for each tract we modeled a main effect of SRT predicting diffusion, and a second model in which the interaction of SRT and NodeID was tested. Using ANOVA, we tested whether including the interaction explained significantly more variance (deviance) than the model with only the main effect of SRT. Table 1 shows that for all tracts, including the interaction term explains significantly more variance (all *F* tests comparing the two models were significant at *p* < .001). This means that associations between SRT and diffusion properties differed in magnitude depending on placement along the tract profile. Significance tests for age and ADHD symptomology covariates are included in Supplemental Table 1.

**TABLE 1.** Behavioral Measures Applied for ADHD Diagnosis.

| Measure | Source | Description of Measure |
| --- | --- | --- |
| Computerized-Diagnostic Interview Schedule for Children (C-DISC) | Shaffer et al., 2000 | The C-DISC is a highly structured, computerized diagnostic interview which can be used to assess over 30 different psychiatric disorders, including ADHD. It consists of a series of questions which, after a minimal training period, can be administered by "lay" interviewers, such as teachers and parents. |
| Disruptive Behavior Disorders Rating Scale (DBD) | Pelham et al., 1992; Graziano et al., 2022 | The DBD assesses symptoms of ADHD, ODD, and CD on a four-point scale regarding the frequency of occurrence. For this study, we controlled for hyperactivity/impulsivity and inattention symptoms measured by the DBD to account for ADHD symptomatology. |
| Impairment Rating Scale (IRS) | Fabiano et al., 2006 | The IRS assesses functional impairment in several key domains and can effectively discriminate between children with and without ADHD. Academic, behavioral, and social impairments are measured by a score of three or higher on a seven-point scale. |

Dual Ph.D. level clinician review was used to determine diagnosis and eligibility for the sample through assessment of three different measures: the Computerized-Diagnostic Interview Schedule for Children (C-DISC), Disruptive Behavior Disorders Rating Scale (DBD), and Impairment Rating Scale (IRS).

With these interaction effects established, we conducted difference tests (essentially independent samples *t*-tests) at each node, assessing high vs. low performers (defined by the median split). Because this amounts to 100 tests for each tract, we corrected for multiple comparisons using the false discovery rate (FDR; (Benjamini & Hochberg, 1995)) procedure. These tract profiles are shown in Figures 9 and 11 (for cortical association pathways) and Figure 10 and 12 (for cerebellar pathways). In the figures, blue marks Node IDs that survived the FDR correction. In general, the results were similar for RNT and HNT, just in opposite directions.

**FIGURE 9.**
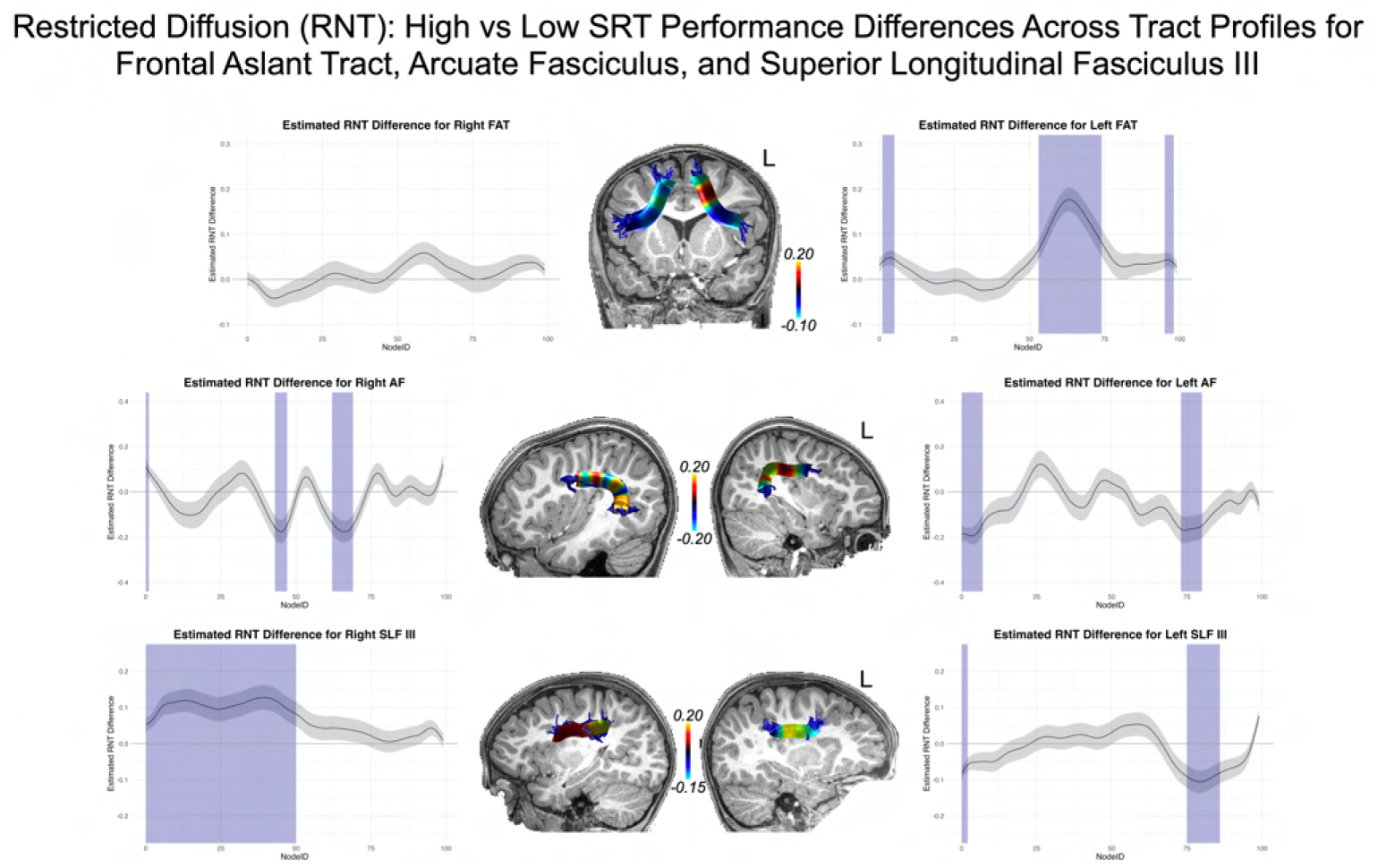
High vs Low Syllable Repetition Task performance differences across tract profiles for Frontal Aslant Tract (FAT), Arcuate Fasciculus (AF), and Superior Longitudinal Fasciculus III (SLF III), for the Restricted Diffusion (RNT) metric. The y-axis reports the difference score for high versus low performers determined by median split. The x-axis reports the Node Index for the 100 nodes along the tract profile. Blue indicates the difference was statistically significant after false discovery rate (FDR) correction. The difference scores from the statistical analysis are mapped to representative tracts overlaid on T1 images for each tract.

**FIGURE 10.**
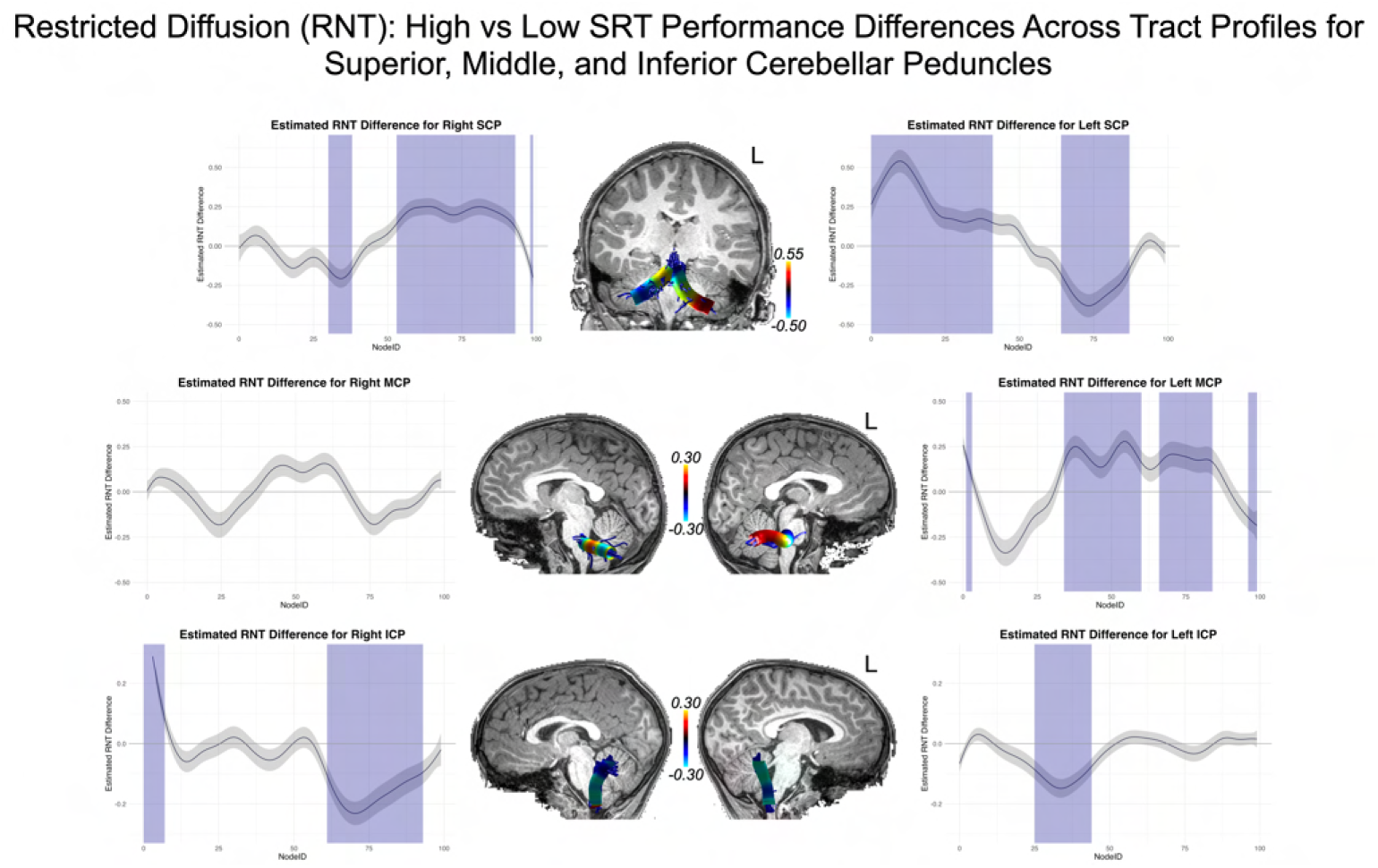
High vs Low Syllable Repetition Task performance differences across tract profiles for Superior (SCP), Middle (MCP), and Inferior (ICP) Cerebellar Peduncles, for the Restricted Diffusion (RNT) metric. The y-axis reports the difference score for high versus low performers determined by median split. The x-axis reports the Node Index for the 100 nodes along the tract profile. Blue indicates the difference was statistically significant after false discovery rate (FDR) correction. The difference scores from the statistical analysis are mapped to representative tracts overlaid on T1 images for each tract.

**FIGURE 11.**
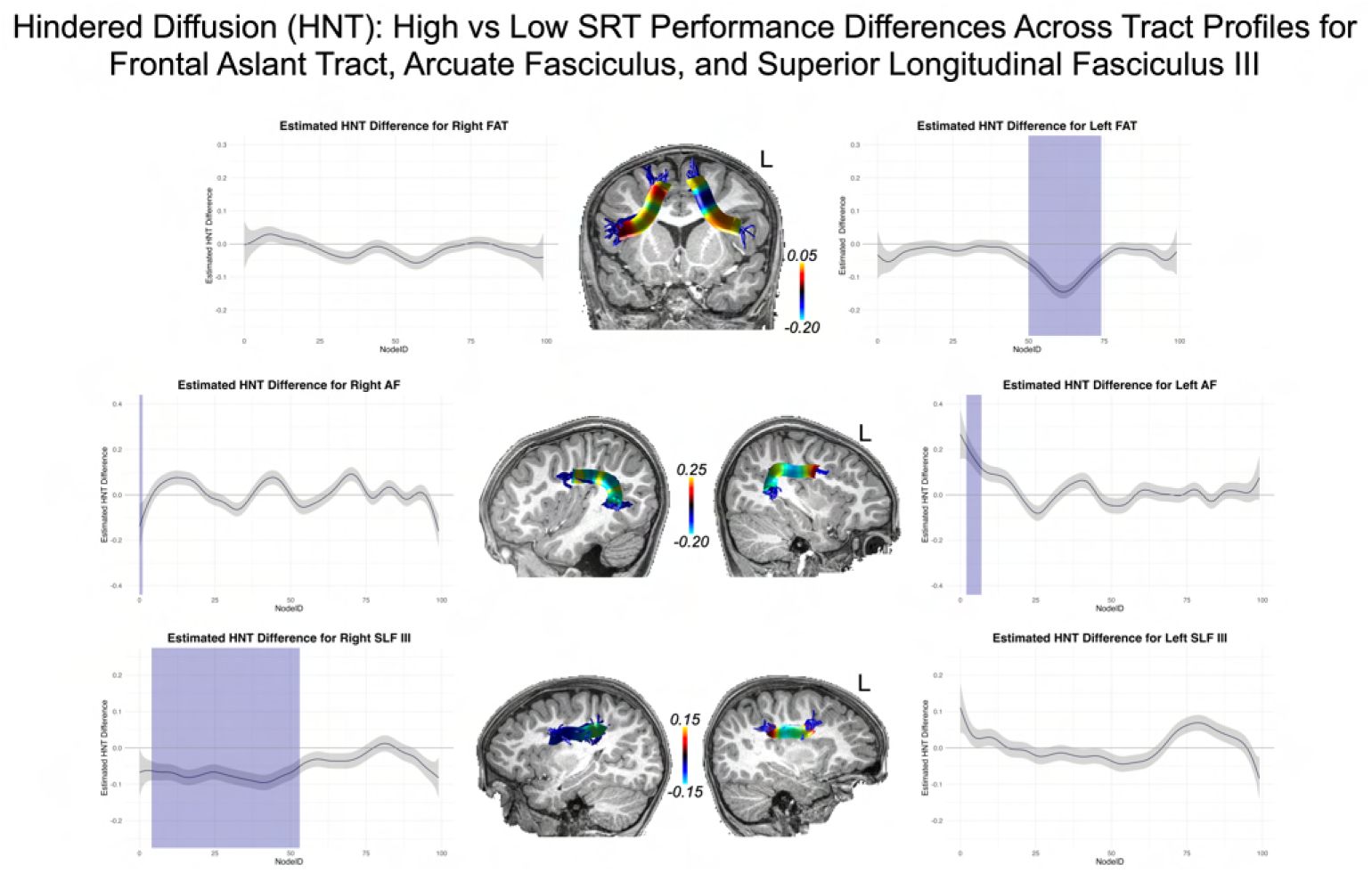
High vs Low Syllable Repetition Task performance differences across tract profiles for Frontal Aslant Tract (FAT), Arcuate Fasciculus (AF), and Superior Longitudinal Fasciculus III (SLF III), for the Hindered Diffusion (HNT) metric. The y-axis reports the difference score for high versus low performers determined by median split. The x-axis reports the Node Index for the 100 nodes along the tract profile. Blue indicates the difference was statistically significant after false discovery rate (FDR) correction. The difference scores from the statistical analysis are mapped to representative tracts overlaid on T1 images for each tract.

**FIGURE 12.**
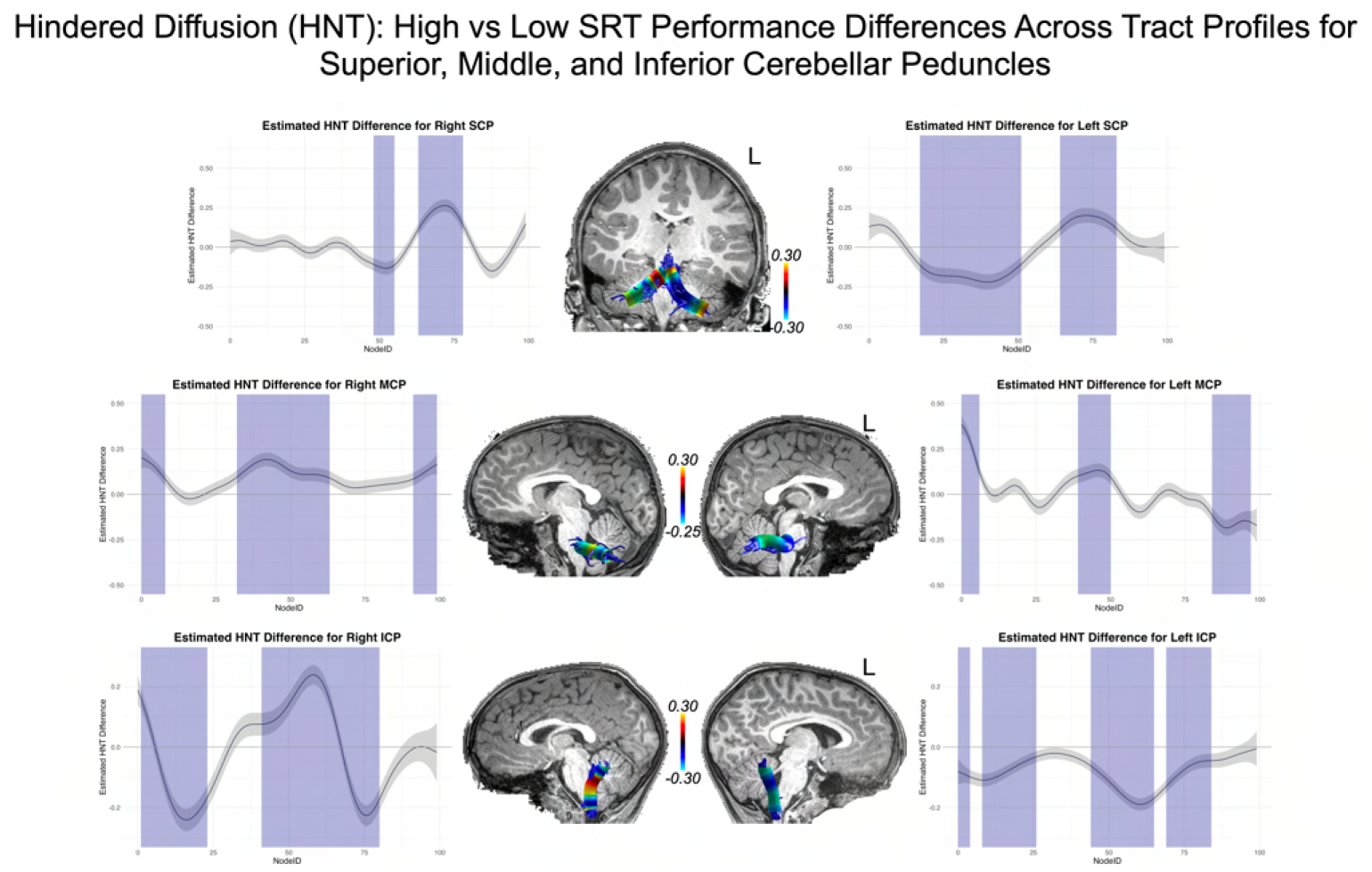
High vs Low Syllable Repetition Task performance differences across tract profiles for Superior (SCP), Middle (MCP), and Inferior (ICP) Cerebellar Peduncles, for the Hindered Diffusion (HNT) metric. The y-axis reports the difference score for high versus low performers determined by median split. The x-axis reports the Node Index for the 100 nodes along the tract profile. Blue indicates the difference was statistically significant after false discovery rate (FDR) correction. The difference scores from the statistical analysis are mapped to representative tracts overlaid on T1 images for each tract.

As the Figures show, for FAT significant nodes were identified along the tract and highlighted in blue. No significant nodes were identified for the right FAT.

Although no significant clusters were identified in AF and SLF III in the whole brain analysis, fiber tract segmentation revealed significant RNT differences between Above and Below Median SRT performance groups at considerable intervals along the tract profiles for SLF III, especially on the right hemisphere. In fact, no differences were found in left SLF III for the HNT metric. There were also significant differences in SRT performance for the left and right AF tract profiles, but these were not extensive, and were at isolated points along the tract.

We also examined significant differences in RNT between SRT performance groups for the cerebellar peduncles. The bilateral SCP and ICP both contained large intervals with significant differences at those nodes. For MCP, differences were only found on the left (for RNT), but bilaterally for HNT.

## 5| DISCUSSION

Updated models of speech implicate a network of grey matter regions and the white matter pathways that connect them in speech production. Despite progress in identifying these regions and pathways, there are many questions that remain about how this speech network develops in early childhood. We examined this development in 4-7-year-olds (some diagnosed with ADHD) who varied considerably in speech ability as measured by a speech task, the SRT. Our results revealed several key findings: 1) performance on the SRT is associated with differences in grey matter cellularity in regions identified as critical for speech production; 2) performance on the SRT further predicts differences in diffusion metrics along white matter tracts related to speech production; 3) the association between speech task performance and grey and white matter cellularity changes as a function of age.

### 5.1| Performance on the SRT is associated with cellularity differences in frontal brain regions and pathways associated with speech

In the whole-brain analysis, we observed that 1) higher RNT in bilateral *pars opercularis* was associated with better performance on the SRT, and 2) reduced RNT and increased HNT in the right pre-SMA/SMA were similarly linked to better SRT performance. However, neither finding survived multiple comparison correction. While these regions align with prior literature, these results should be interpreted cautiously. We provide a brief discussion of these findings with this limitation in mind, starting with the *pars opercularis* before addressing the pre-SMA/SMA.

The left *pars opercularis* has been implicated in speech neurobiology models. For instance, in GODIVA (Guenther, 2016), this region may house cell populations representing syllable motor programs independent of semantic content (Ghosh et al., 2008). Similarly, Hickok’s model (Hickok et al., 2023) posits that the *pars opercularis* is involved in syllable sequencing. While our findings align with these models, the bilateral association observed in our data may reflect developmental factors, and previous research suggests that lateralization of language function increases with age. For example, Holland et al. (S. K. Holland et al., 2001) showed that left lateralization in the inferior frontal cortex strengthens from ages 7-to-18 during a verb generation task. Similarly, Olulade et al. (Olulade et al., 2020) found that younger children (aged 4-8) show stronger activation in homologues of the right hemisphere during fMRI language tasks, with this pattern decreasing through adolescence. Our bilateral findings in *pars opercularis* may therefore reflect the substantial contribution of the right hemisphere to speech in younger children. Furthermore, age moderated the association between SRT performance and RNT in the left *pars opercularis*, with stronger associations observed in older children, consistent with increased left lateralization over development.

**TABLE 2.** Model Fit Indices and Comparisons for Each Tract Profile.

| Tract | F (p-value) | Main Effect AIC | % Deviance Explained (ME) | Interaction AIC | % Deviance Explained (Int) |
| --- | --- | --- | --- | --- | --- |
| Left FAT | 9.1 *** | -33293.9 | 84.4 | -33815.3 | 85.4 |
| Right FAT | 8.8 *** | -34275.3 | 85.9 | -34756.8 | 86.7 |
| Left AF | 6.8 *** | -17568.4 | 43.7 | -17878.1 | 46.4 |
| Right AF | 9.2 *** | -16672.4 | 31.2 | -17138.4 | 35.2 |
| Left SLF III | 8.7 *** | -31870.7 | 70.8 | -32314.8 | 72.5 |
| Right SLF III | 7.9 *** | -30573.8 | 68.7 | -30937.1 | 70.2 |
| Left SCP | 9.7 *** | -3850.8 | 13.7 | -4296.7 | 21.2 |
| Right SCP | 8.91 *** | -6406.8 | 14.0 | -6827.3 | 20.8 |
| Left MCP | 7.07 *** | -7257.1 | 25.7 | -7539.2 | 28.3 |
| Right MCP | 7.37 *** | -7516.1 | 28.8 | -7821.1 | 31.5 |
| Left ICP | 12.4 *** | -27445.7 | 53.2 | -28132.1 | 57.0 |
| Right ICP | 16.8 *** | -14909.4 | 24.6 | -15979.6 | 33.3 |

*Note.* \*\*\* = $p < .001$ ; AIC = Akaike Information Criterion; ME = Main Effect, Int = Interaction Effect. FAT = Frontal Aslant Tract. AF = Arcuate Fasciculus; SLF III = Superior Longitudinal Fasciculus III; SCP, MCP, ICP = Superior, Middle, and Inferior Cerebellar Peduncles.

The relationship between restricted diffusion and performance remains speculative, but can be informed by prior histologic studies. Palmer et al. (Palmer et al., 2022) suggested that age-related increases in restricted diffusion in grey matter could reflect changes such as increased myelination, neurite density, dendritic sprouting, or cell body growth (neuronal or glial). White et al. (White, Leergaard, et al., 2013a; White, McDonald, Farid, et al., 2013; White et al., 2014), in developing the RSI model, highlighted its sensitivity to these microstructural properties. Based on this, we cautiously suggest that the observed associations may reflect these underlying tissue changes. However, given that these findings did not survive the cluster correction, they should be interpreted with appropriate caution and considered preliminary.

In the white matter of FAT, we can speculate that these associations have more to do with myelination of the pathway, which reduces the volume of extracellular space and also reduces the permeability of axonal membranes. This would lead to an increase in the signal in the restricted compartment. The AFQ analysis of the FAT showed that this effect was predominantly unilateral. In fact, no performance differences in the right FAT were statistically significant after FDR correction. Thus, unlike the finding in *pars opercularis* grey matter, the finding in FAT was prominently left lateralized. Moderating effects of age were also left lateralized. It is possible that for FAT white matter, connecting inferior frontal gyrus and pre-SMA/SMA cortical regions, the left FAT is more critical for speech than the right FAT (Dick et al., 2019). This is suggested by studies in adults, which have shown that left FAT integrity is associated with stuttering (Kemerdere et al., 2016; Kronfeld-Duenias et al., 2016; Misaghi et al., 2018), speech arrest during tumor resection and intraopertive electrostimulation (Fujii et al., 2015; Vassal et al., 2014), post-stroke aphasia (Basilakos et al., 2014) and speech apraxia (Zhong et al., 2022). In this latter study, damage to the white matter and not the grey matter was the best predictor of speech impairment, suggesting that disconnection of the connectivity is a stronger factor for predicting motor speech deficits. Our findings are consistent with this possibility.

We also identified a small cluster in the right pre-SMA/SMA showing an association between SRT performance and reduced RNT/increased HNT. However, this cluster was small and did not survive cluster correction, so these findings should be interpreted with caution. The involvement of this region is consistent with previous evidence on the role of bilateral pre-SMA/SMA in speech production (Alario et al., 2006; Bohland & Guenther, 2006; Tremblay & Gracco, 2009, 2010). For example, stimulation of pre-SMA / SMA during awake surgery can cause speech arrest (Lu et al., 2021), and Bohland and Guenther (Bohland & Guenther, 2006) showed that even simple syllable sequences engage the bilateral pre-SMA/SMA as part of the basic speech production network.

The direction of the effect—less restricted diffusion (and more hindered diffusion) being associated with better performance—differs from our other findings, making interpretation more challenging. Changes in diffusion properties can reflect various microstructural processes, such as increased dendritic pruning, which might enhance the processing efficiency of neural assemblies in the pre-SMA/SMA and result in reduced restricted diffusion and increased hindered diffusion signals. While this is a plausible explanation, it remains speculative and would require histological validation.

### 5.2| Performance on the SRT is associated with cellularity differences in cerebellar white and grey matter

The whole brain analysis revealed that 1) higher RNT and HNT in bilateral Crus I and II was associated with better SRT performance; and 2) higher RNT and HNT was associated with cerebellar white matter pathways: the SCP (for RNT only), MCP, and ICP. While for RNT, findings from cerebellar grey matter and MCP/ICP survived cluster correction, the SCP clusters did not (no clusters survived for HNT). Thus, associations with SCP should be interpreted with this caveat in mind.

The role of the cerebellum in speech motor control has long been attributed to the organization and coordination of motor commands necessary for speech production, but is particularly important for rapid speech (Ackermann, 2008; Guenther, 2016) or for more complex syllable sequences (Bohland & Guenther, 2006). The developmental importance of finely timed speech has been demonstrated in studies of children with early speech deficits, which found that poorer speech outcomes were associated with right cerebellar tumors (A. Morgan et al., 2011). This is consistent with adult cerebellar lesion studies (Urban et al., 2003), and the established connectivity of inferior frontal regions with the contralateral right cerebellum (A. Morgan et al., 2011). Results from the present study indicated bilateral findings in the grey matter of the cerebellum, especially in the Crus I and Crus II cerebellar lobules. However, fMRI activation during speech and foci collected from meta-analyses of word and sentence reading have highlighted bilateral associations between speech and cerebellar grey matter (Brown et al., 2005; Guenther, 2016; Turkeltaub et al., 2002). Further, bilateral activation of Crus I and Crus II during speech production tasks has been reported (Bohland & Guenther, 2006; Correia et al., 2020; Geva et al., 2021; Ghosh et al., 2008; Peeva et al., 2010; Shuster & Lemieux, 2005). Thus, our findings are in line with previous literature that suggests motor control of speech is not limited to the right cerebellum.

The connectivity of the cerebellum with cortical and subcortical structures in the cerebral cortex is established through the cerebellar peduncles, bilateral white matter tracts that have been shown to play a role in aspects of speech fluency and motor speech production (Guenther, 2016; Johnson et al., 2022; Jossinger et al., 2023). Jossinger and colleagues investigated the role of the cerebellar peduncles in different components of speech processing, and found that verbal fluency was associated with the right SCP, whereas speaking rate was associated with the right MCP (Jossinger et al., 2023). A similar pattern is observed in children who stutter, as early developmental differences have been demonstrated between children who stutter and their age-matched peers in the right ICP (Johnson et al., 2022). However, the association between the cerebellar peduncles and motor speech production is not restricted to the right cerebellar peduncles. In adults who stutter, speech rate has been associated with white matter cellular properties in the left ICP (Jossinger et al., 2021). In addition, reduced integrity of the white matter has been bilaterally observed for all three cerebellar peduncles in adolescents and adults who stutter (Connally et al., 2014), confirming the notion that motor speech production is necessarily lateralized right. Our bilateral findings in the SCP, MCP, and ICP support the idea that key aspects of speech production may be organized bilaterally within the white matter pathways that connect the cerebellum to the cerebral cortex. The findings in the SCP from the whole brain analysis did not survive cluster correction, but our analysis of the SCP tract profiles revealed several significant nodes, which suggests that there may be some relationship between white matter cellular properties and speech performance that requires additional study to tease apart. Although both the left and right ICPs showed significant nodes, only the left MCP demonstrated a difference in SRT performance between nodes. We may have observed these differences because the cerebellar peduncles play different roles in motor speech processing, as noted by Jossinger et al (Jossinger et al., 2023), but understanding this relationship requires further analysis of different speech properties.

The findings from the present study demonstrate that greater differences in performance on a speech production task is associated with higher RNT and HNT in grey matter of the cerebellum, and higher RNT and lower HNT in white matter of the cerebellar peduncles (based on the tract profile analysis). Based on our knowledge of how RSI modulates developmental and maturational processes, we can speculate on the nature of the association, but we are unable to make definitive claims about the relationship between the restricted diffusion signal and speech performance without using more direct measurements. Developmental processes such as dendritic sprouting, myelination, and increasing neurite diameter result in a decrease in extracellular space, which in turn causes the signal to increase within the cellular compartment (Palmer et al., 2022). In particular, higher myelination is associated with greater speed of conductance for information traveling between neurons (Jossinger et al., 2021; Palmer et al., 2022; Zhong et al., 2022). We may tentatively conclude that an increase in RNT and decrease in HNT, possibly driven by developmental processes such as myelination, could lead to faster conduction of information and thus better performance on a speech task. This would be in line with the results from our whole brain analysis, which suggested that higher restricted diffusion signal was positively associated with speech performance, and that age moderated the strength of this association, which increased with age. While these conclusions are speculative, understanding the relationship between microstructural properties of grey and white matter and speech performance allows us to establish a relationship that may be further investigated in the future.

### 5.3| Tract profile analysis reveals more detailed relationship between SRT performance and dorsal stream fiber pathways

Although the whole-brain analysis did not reveal significant associations between SRT performance and the AF or SLF III, the AFQ analysis identified significant differences in RNT between the Above Median and Below Median SRT groups, albeit with minimal differences for the AF. Both the AF and SLF III are known to connect regions critical for motor speech function, such as the IFG with STS/STG, and SMG (Bernal & Ardila, 2009; Duffau et al., 2003; Hickok & Poeppel, 2004, 2007). These findings, though subtle, may provide converging evidence for the role of these tracts in motor speech function.

It is important to interpret these results cautiously. GAMs are highly sensitive to small differences, and the whole-brain analysis did not reveal associations between RNT and SRT performance in the AF or SLF III. However, tractography at the individual participant level leverages the native MRI space of each individual, whereas whole-brain analyses rely on registration to a template space, which can reduce spatial specificity and sensitivity for structures of interest. Traditional fiber tracking methods, which average diffusion metrics across an entire fiber bundle, may miss localized relationships within the tract. Diffusion metrics can vary significantly along the trajectory of a tract (Yeatman et al., 2011). By creating tract profiles that examine diffusion metrics at specific nodes along the tract, methods like AFQ can reveal relationships that might otherwise remain obscured. Investigating white matter tracts with such approaches is therefore a crucial step toward a more nuanced understanding of the relationship between white matter microstructural properties and diffusion metrics.

Our results underscore the potential value of the AFQ findings in advancing our understanding of the neural speech network. This is especially the case for right SLFIII, which showed a sustained difference across the profile of a large part of the anterior tract. It was also evident for left SLFIII in the posterior part of the tract in the white matter near the supramarginal gyrus. This is consistent with Duffau’s work showing the importance of this white matter for processing phonological information during speech production (Duffau et al., 2003). In that study, electrostimulation during awake surgery of white matter connecting SMG and inferior frontal gyrus disrupted speech production. Thus, our finding showing such an association in a developmental sample is novel and provides additional support for the importance of this pathway to speech production.

### 5.4| Limitations

One limitation of the current study regards the sample makeup, which included both children who are typically developing and children with ADHD, and contained a higher percentage of boys than girls. Children who are atypically developing, such as those with ADHD, are at risk for developing comorbid psychiatric and speech disorders (Booster et al., 2012), and a mixed sample over-represented by boys may not be generalizable to a broader population. However, few studies focusing on the development of neural speech networks in early childhood exist, and the findings from this study provide critical background for understanding the implementation of speech production in the brain at early stages of speech development. We did not find performance differences on the SRT, and we further controlled for ADHD symptomology. This suggests that treating the ADHD and TD children as a representative sample for performance on this task is justified.

A second limitation concerns the speech task implemented in the current research, the SRT. For this study, we focused on a single speech task that measured the repetition of multisyllabic utterances. Although the SRT measures expressive speech production (Shriberg et al., 2009), it is not sensitive to the different phases of speech that may occur during repetition. For example, the SRT is unable to distinguish between prearticulatory encoding, and the actual transformation of phonological planning into motor execution of speech. The SRT is also a speech task that measures the repetition of single, multisyllabic nonwords, and thus cannot provide insight into how connected speech may play a role in speech processing. Despite these limitations, the SRT is a child-friendly phonemic task that has been validated as an assessment of motor planning difficulties (Rvachew & Matthews, 2017), and may be considered as part of a battery of speech and language tasks in the future to provide a more complete picture of speech development.

Finally, we must carefully consider what our diffusion metrics of interest actually measure. Although we can provide speculation for what cellular processes are modulated by restricted and hindered diffusion, we cannot definitively identify the developmental and maturational processes underlying differences in diffusion metrics. The issue of specificity is a problem for diffusion measures more broadly, not just RSI. In fact, a major strength of RSI beyond traditional diffusion measures, such as diffusion tensor imaging (DTI), is that we are able to observe differences in cellularity in both grey and white matter. While RSI has its constraints, much like other DWI reconstruction methods, it provides a more complete picture of neural development across grey matter regions of interest and white matter pathways.

## 6| CONCLUSION

Characterizing the structural development of the neural speech network in early childhood is important for understanding speech acquisition. In this investigation, we found that 1) performance on the SRT is associated with differences in grey matter cellularity in regions identified as critical for speech production, including inferior frontal gyrus and pre-SMA/SMA and in cerebellum. An important caveat is that the cortical findings do not survive cluster correction for multiple comparisons; 2) performance on the SRT further predicts differences in diffusion metrics along white matter tracts related to speech production, especially left FAT, left and right SLFIII, and the cerebellar peduncles; 3) the association between speech task performance and grey and white matter cellularity changes as a function of age in these regions and pathways. The findings suggest that individual differences in speech performance are reflected in structural grey and white matter differences as measured by restricted and hindered diffusion properties, and offer important insights into how the neural speech network develops in children during early childhood.

## 6.2| Acknowledgments

We thank the parents and children who participated in the AHEAD study from which these data were analyzed. We also thank Anders Dale and Donald Hagler for help with RSI reconstruction, and M. Okan Irfanoglu and Carlo Pierpaoli for recommended DWI reconstruction steps for ABCD diffusion acquisition protocols, and improvements to those protocols.

## Abbreviations

ADHD: Attention-Deficit/Hyperactivity Disorder

