## Supplemental Tables for "Structural Development of Speech Networks in Young Children"

### 0.1 | Supplementary Tables

#### Effects of Age and ADHD Symptomology for HNT

| Tract | Coefficient | Estimate (Std Error) | t-value |
|---|---|---|---|
| Left Frontal Aslant Tract | Age | -0.021 (0.003) | -6.776*** |
| | Mean Hyperactivity | 0.065 (0.005) | 13.386*** |
| | Mean Inattention | -0.038 (0.004) | -8.606*** |
| Right Frontal Aslant Tract | Age | -0.046 (0.003) | -15.097*** |
| | Mean Hyperactivity | 0.027 (0.005) | 5.217*** |
| | Mean Inattention | 0.014 (0.005) | 3.142** |
| Left Arcuate Fasciculus | Age | -0.033 (0.003) | -9.480*** |
| | Mean Hyperactivity | -0.046 (0.006) | -7.987*** |
| | Mean Inattention | 0.0216 (0.005) | 4.251*** |
| Right Arcuate Fasciculus | Age | -0.030 (0.003) | -10.041*** |
| | Mean Hyperactivity | 0.008 (0.005) | 1.507 |
| | Mean Inattention | -0.022 (0.004) | -5.058*** |
| Left Superior Longitudinal Fasciculus III | Age | -0.013 (0.003) | -3.950*** |
| | Mean Hyperactivity | 0.067 (0.006) | 11.937*** |
| | Mean Inattention | -0.044 (0.005) | -8.803*** |
| Right Superior Longitudinal Fasciculus III | Age | -0.035 (0.003) | -10.695*** |
| | Mean Hyperactivity | 0.042 (0.006) | 7.628*** |
| | Mean Inattention | -0.014 (0.005) | -2.966** |
| Left Superior Cerebellar Peduncle | Age | -0.042 (0.012) | -3.633*** |
| | Mean Hyperactivity | 0.109 (0.021) | 5.306*** |
| | Mean Inattention | -0.041 (0.020) | -2.008* |
| Right Superior Cerebellar Peduncle | Age | 0.0154 (0.009) | 1.686 |
| | Mean Hyperactivity | 0.002 (0.014) | 0.169 |
| | Mean Inattention | -0.004 (0.013) | -0.308 |
| Left Middle Cerebellar Peduncle | Age | -0.034 (0.007) | -4.815*** |
| | Mean Hyperactivity | -0.070 (0.012) | -5.923*** |
| | Mean Inattention | 0.053 (0.010) | 5.008*** |
| Right Middle Cerebellar Peduncle | Age | -0.013 (0.007) | -1.930 |
| | Mean Hyperactivity | -0.030 (0.011) | -2.591** |
| | Mean Inattention | 0.008 (0.010) | 0.807 |
| Left Inferior Cerebellar Peduncle | Age | -0.020 (0.005) | -4.383*** |
| | Mean Hyperactivity | 0.032 (0.008) | 4.216*** |
| | Mean Inattention | 0.036 (0.007) | 5.373*** |
| Right Inferior Cerebellar Peduncle | Age | 0.022 (0.008) | 2.764** |
| | Mean Hyperactivity | 0.002 (0.012) | 0.139 |
| | Mean Inattention | -0.033 (0.011) | -2.983** |

#### Effects of Age and ADHD Symptomology for RNT

| Tract | Coefficient | Estimate (Std Error) | t-value |
|---|---|---|---|
| Left Frontal Aslant Tract | Age | 0.035 (0.003) | 10.993*** |
| | Mean Hyperactivity | -0.062 (0.005) | -11.922*** |
| | Mean Inattention | 0.033 (0.005) | 6.922*** |
| Right Frontal Aslant Tract | Age | 0.041 (0.003) | 13.456*** |
| | Mean Hyperactivity | -0.041 (0.005) | -8.060*** |
| | Mean Inattention | -0.001 (0.005) | -0.267 |
| Left Arcuate Fasciculus | Age | 0.039 (0.004) | 8.873*** |
| | Mean Hyperactivity | 0.029 (0.007) | 4.035*** |
| | Mean Inattention | -0.014 (0.006) | -2.215* |
| Right Arcuate Fasciculus | Age | 0.021 (0.004) | 5.061*** |
| | Mean Hyperactivity | -0.081 (0.007) | -11.745*** |
| | Mean Inattention | 0.074 (0.006) | 11.911*** |
| Left Superior Longitudinal Fasciculus III | Age | 0.005 (0.004) | 1.278 |
| | Mean Hyperactivity | -0.092 (0.007) | -13.995*** |
| | Mean Inattention | 0.049 (0.006) | 8.287*** |
| Right Superior Longitudinal Fasciculus III | Age | 0.069 (0.004) | 15.978*** |
| | Mean Hyperactivity | -0.105 (0.007) | -15.187*** |
| | Mean Inattention | 0.059 (0.006) | 9.512*** |
| Left Superior Cerebellar Peduncle | Age | 0.024 (0.013) | 1.887 |
| | Mean Hyperactivity | -0.001 (0.022) | -0.005 |
| | Mean Inattention | 0.012 (0.022) | 0.562 |
| Right Superior Cerebellar Peduncle | Age | 0.099 (0.012) | 8.126*** |
| | Mean Hyperactivity | 0.04 (0.022) | 1.793 |
| | Mean Inattention | 0.063 (0.017) | 3.733*** |
| Left Middle Cerebellar Peduncle | Age | -0.022 (0.010) | -2.205* |
| | Mean Hyperactivity | -0.095 (0.016) | -5.822*** |
| | Mean Inattention | 0.119 (0.015) | 7.802*** |
| Right Middle Cerebellar Peduncle | Age | 0.095 (0.010) | 9.786*** |
| | Mean Hyperactivity | -0.003 (0.015) | -0.188 |
| | Mean Inattention | -0.010 (0.014) | -0.686 |
| Left Inferior Cerebellar Peduncle | Age | 0.013 (0.004) | 2.871** |
| | Mean Hyperactivity | -0.003 (0.007) | -0.371 |
| | Mean Inattention | -0.005 (0.006) | -0.805 |
| Right Inferior Cerebellar Peduncle | Age | 0.033 (0.007) | 4.896*** |
| | Mean Hyperactivity | -0.055 (0.010) | -5.390*** |
| | Mean Inattention | 0.050 (0.010) | 5.088*** |

**TABLE 1** Table 1 provides an overview of the effects of age and ADHD symtomology for the hindered (HNT) and restricted (RNT) restriction spectrum imaging (RSI) components. Across most of the tracts, age, mean hyperactivity, and mean impulsivity were significant for both HNT and RNT. Notably, these coefficients were not significant in the Right Superior Cerebellar Peduncle (SCP) for HNT, and the Left SCP for RNT. * $p$ < .05. ** $p$ < .01. *** $p$ < .001
